# Comparison and benchmark of deep learning methods for non-coding RNA classification

**DOI:** 10.1101/2023.11.24.568536

**Authors:** Constance Creux, Farida Zehraoui, François Radvanyi, Fariza Tahi

**Affiliations:** Université Paris-Saclay, Univ Evry, IBISC, 91020 Evry-Courcouronnes, France; Molecular Oncology, PSL Research University, CNRS, UMR 144, Institut Curie, Paris, France

## Abstract

The involvement of non-coding RNAs in biological processes and diseases has made the exploration of their functions crucial. Most non-coding RNAs have yet to be studied, creating the need for methods that can rapidly classify large sets of non-coding RNAs into functional groups, or classes. In recent years, the success of deep learning in various domains led to its application to non-coding RNA classification. Multiple novel architectures have been developed, but these advancements are not covered by current literature reviews. We present an exhaustive comparison of the different methods proposed in the state-of-the-art and describe their associated datasets. Moreover, the literature lacks objective benchmarks. We perform experiments to fairly evaluate the performance of various tools for non-coding RNA classification on popular datasets. The robustness of methods to non-functional sequences and sequence boundary noise is explored. We also measure computation time and CO_2_ emissions. With regard to these results, we assess the relevance of the different architectural choices and provide recommendations to consider in future methods. Datasets and reproducible codes are available at https://evryrna.ibisc.univ-evry.fr/evryrna/ncBench.

**Author summary:** RNA can either encode proteins, which perform different functions in the genome, or be non-coding. Non-coding RNAs represent around 98% of the genome, and were long thought to be non-functional. It has now been proven that non-coding RNAs can have diverse biological functions and be involved in diseases. A large proportion of non-coding RNAs has not yet been studied. The function of specific non-coding RNAs can be studied experimentally, but experiments are costly and time-consuming. One possibility to massively characterize the function of non-coding RNAs is to use computational methods to classify them into functional groups, or classes. Recent computational methods for non-coding RNA classification are all based on deep learning, as it leads to faster runtime and improved performance. Our work presents and compares the different approaches adopted in the state-of-the-art, as well as the non-coding RNA datasets that are used. We also present a comprehensive benchmark, measuring classification performance in different conditions, computation time, and CO_2_ emissions. The descriptions and comparisons provided are meant to guide researchers in the field, whether wanting to use existing tools or to develop new ones.

## Introduction

The definition of non-coding RNA (ncRNA) classes of similar functions started in the 1950s, with the discovery of tRNAs (transfer RNAs) and rRNAs (ribosomal RNAs) [1]. The description of other ncRNA classes involved in cell maintenance followed, but research still mainly focused on proteins or protein-coding genes. It is in the 2000s, when efforts like the Human Genome Project [2] highlighted that 98% of the genome is non-coding, that attention started to turn to ncRNA. Novel classes of ncRNAs with regulatory functions, like miRNA (microRNA) or lncRNA (long ncRNA), were described. Studies have shown that ncRNAs are implicated in various biological processes and diseases [3–5], underscoring their potential as biomarkers and therapeutic targets, and thus showing that studying their functions is important. Knowing their precise functions requires specific experimental studies, which are resource-intensive. To focus these resources on ncRNAs of interest, a first functional characterization can be done computationally on a large scale. Groups can be formed to gather ncRNAs that perform the same function. This is typically done at two levels. The first is ncRNA families, which are quite specific, and represent ncRNAs that share the same function, have similar structural motifs, and have sequence homology. The second is ncRNA classes, which are broader. They also represent ncRNAs sharing a function and structural motifs, but sequence homology is no longer a constraint. In this work, we focus on ncRNA class prediction. Still, it is worth noting that ncRNA family prediction has also garnered interest in the community and has been addressed in multiple works [6–8].

Many ncRNA classes have been identified over the years. It is typical to separate them based on whether they perform ‘housekeeping’ functions, i.e., they are involved in cell viability, or whether they are ‘regulatory’, i.e., they regulate gene expression at the epigenetic, transcriptional or post-transcriptional level. We represent in Fig 1 these two types of functions, naming some ncRNA classes that are commonly included in ncRNA classification problems. Each of these classes is associated to a specific function. For example, tRNAs [9] have a distinctive cloverleaf-like structure, and their function is to carry amino acids to the ribosome for protein synthesis. MiRNAs [10] range from 20 to 26 nt long, their precursors have a hairpin structure, and they are molecules that bind mRNA to regulate expression. They typically repress the expression of downstream genes, and can sometimes even silence it completely. Some classes are better known than others, and new classes could still be discovered as new functions of ncRNAs are uncovered: to this day, there is no fixed set of ncRNA classes. LncRNAs particularly suffer from this lack of definition [11]. RNAs belonging to this class are disparate and can have multiple functions. They are mostly defined by their length over 200 nt and their position in the genome (antisense, intronic, …). Due to this lack of precision, ncRNA classification problems tend to focus on classes of small ncRNAs (sncRNAs).

**Fig 1.**
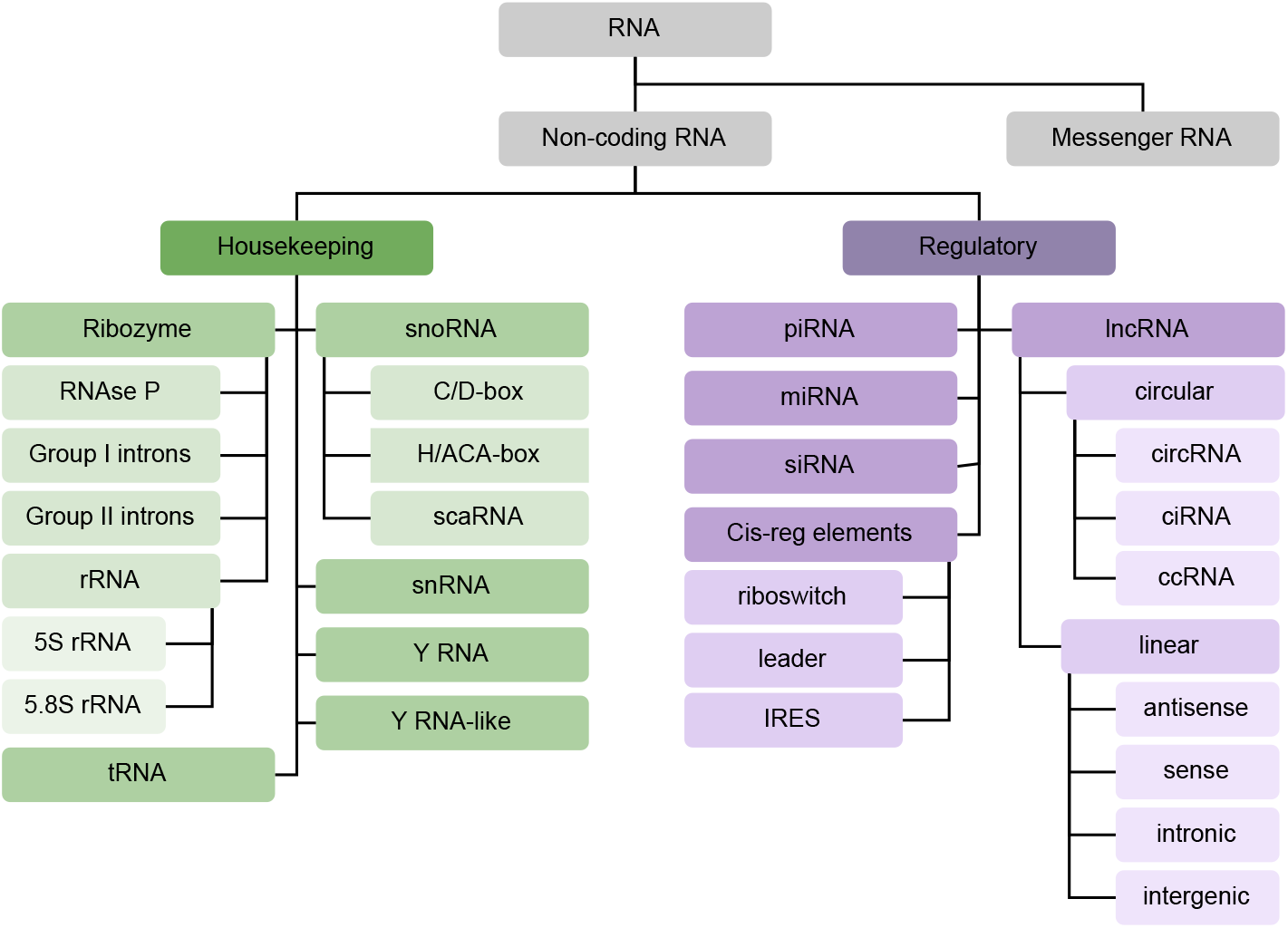
Taxonomy of ncRNA classes. Non-coding RNA functions can be separated into housekeeping (green) and regulatory (purple). Classes can be subdivided depending on the level of description. The classes represented are examples that are often mentioned in the literature, or that are of interest in the datasets mentioned below.

Several tools have been proposed since the 2000s to predict ncRNA classes. The use of computational resources enables the obtention of results faster than experimental methods, as multiple ncRNAs can be studied at the same time. The first approaches relied mainly on alignment algorithms and standard machine learning. In recent years, deep learning (DL) has been applied to ncRNA classification, showing improved performance and an ability to classify even larger ncRNA datasets. DL is now the prevalent approach in the field. The term ‘ncRNA classification’ can have different meanings in the literature. It is sometimes used to refer to the prediction of specific ncRNA classes [12–21]. This problem has been widely covered in reviews [22–25]. It can also refer to multi-class classification, taking as input a set of ncRNAs and associating each one to a ncRNA class, which is the scope of this review.

The literature lacks comparisons of recent multi-class ncRNA classifiers. In an article published in 2017 [26], earlier methods for ncRNA classification are presented. Only one work, published in 2019 [23], covers the use of DL for multi-class ncRNA classification. We count twelve DL-based ncRNA classifiers in the state-of-the-art; eleven of them were presented after its publication and have not been covered in other literature.

This lack of reviews is an issue in the field, as the performance comparisons proposed in publications accompanying the different tools are unreliable. Indeed, comparisons between methods are often made without re-executing tools and instead simply using the scores announced in previous publications. In these instances, results are not always obtained with comparable experimental protocols. Moreover, the main dataset used for evaluation presents a data leakage problem; thus, reported performance is biased.

We propose to fill a gap in the literature by presenting an exhaustive review of the different DL approaches for multi-class ncRNA classification. We describe the choices in DL architecture, and explain the differences between datasets used for evaluation. Moreover, we conduct a fair comparison of performance between current tools. We evaluate global and per-class prediction results. The tools’ ability to recognize non-functional sequences and to disregard sequence boundary noise is reported. We also assess resource intensity by measuring computation time as well as CO_2_ emissions.

This paper is organized as follows: the first section presents the state-of-the-art of DL for ncRNA classification, exhaustively describing current classifiers and the accompanying datasets. In the next section we present the results of our comparison of different tools in terms of global and per-class performance, robustness to noise, as well as computation time and CO_2_ emissions. The section that follows discusses these observations. The Methods section presents the details of our experiments.

### Deep learning for ncRNA classification

#### State-of-the-art classifiers

Earlier methods for ncRNA classification were primarily based on alignment algorithms [6, 27–29] or standard machine learning algorithms [30–32]. Feature extraction in these methods is a particularly crucial step, with a high impact on performance. Current ncRNA classifiers are all based on deep learning: most methods are based on Convolutional Neural Networks (CNNs), Recurrent Neural Networks (RNNs), or a combination of both.

Applications of DL to ncRNA classification started with multiple CNN-based methods: nRC [33], ncRDeep [34], ncrna-deep^1^ [35], RPC-snRC [36], ncRDense [37] and imCnC [38]. nRC pioneered the use of DL in the field. As input, ncRNAs are represented by vectors containing information on the presence in their secondary structure of discriminative sub-structures. Two convolutional layers extract relevant information, and Fully-Connected Layers (FCLs) are used for classification. ncRDeep is another method based on two convolutions followed by FCLs. However, the input is a simple representation of the sequence, a matrix created from one-hot encodings of nucleotides. In the method ncrna-deep, the number of convolutional layers rises to five. Authors tested multiple sequence encodings (space-filling curves or one-hot encoding of k-mers^2^ of length 1, 2 or 3). A one-hot encoding of 1-mers, which corresponds to the same representation as ncRDeep, was selected for its superior results. This representation is also used by RPC-snRC, and fed to three consecutive Dense blocks (based on DenseNets, multiple convolutional layers with dense connections), with a final FCL for classification. ncRDense is an evolution of ncRDeep. In addition to sequence information, the method includes a second input: a matrix with multiple representations of nucleotides and one-hot encoding of the secondary structure. Both inputs are processed with independent DenseNets and merged. A final convolutional layer is followed by FCLs for classification. imCnC stands out as a multi-source CNN. Based on the idea that ncRNAs are related to chromatin states and epigenetic modifications, the approach builds a CNN for each input source (such as the sequence and different epigenetic information), before merging representations to classify samples with FCLs.

RNA sequences can be viewed as sentences, making the use of RNNs relevant to extract meaningful information from ncRNAs, as done by the methods ncRFP [39] and NCYPred [40]. ncRFP uses a bidirectional Long Short-Term Memory RNN (bi-LSTM). When learning a representation for a nucleotide, it is able to take into account close and distant elements, parsing the sequence both forward and backward. An attention mechanism allocates more weight to important information, and FCLs are used for classification. NCYPred also uses a bi-LSTM and FCLs for classification. The main difference between the two is that in ncRFP, each nucleotide is encoded separately, while in NCYPred the sequence is first transformed into overlapping 3-mers, and it is these 3-mers that are encoded.

Several methods combine both approaches, first using RNNs for sequence representation, from which CNNs then extract relevant information. This is the case for ncDLRES [41], ncDENSE [42] and MFPred [43]. ncDLRES is an updated version of ncRFP’s architecture. The bi-LSTM is replaced by a dynamic LSTM, which only parses the sequence in the forward sense, but has the advantage of allowing inputs of varying length. For classification, the FCLs are replaced by a Residual Neural Network (ResNet), which is composed of multiple convolutional layers with skip connections. ncDENSE further evolves that architecture, replacing the dynamic LSTM by a dynamic bidirectional Gated Recurrent Unit (bi-GRU), and the ResNet by a DenseNet. MFPred proposes a rich input representation: sequences are encoded into four sets of features, each processed by a dynamic bi-GRU, reshaped into matrices, and fused. Compared to ncDENSE, the DenseNet is replaced by a Squeeze and Excitation ResNet (SE ResNet), which models interdependencies between the four matrices of learned sequence encodings.

Two existing methods do not fall into the above categorization: RNAGCN [44] and Graph+DL [45], both based on graphs. Taking the secondary structure as input, RNAGCN uses a Graph Neural Network (GNN) with FCL for classification. GNNs are able to learn meaningful representations of non-Euclidean data. RNAGCN is, to date, the only method directly using ncRNA secondary structure without manual feature extraction, removing a source of error. The approach Graph+DL^3^ is based on features extracted from the secondary structure. Authors propose a protocol to obtain a rich graph-theory-based representation of structures. Other than the extensive feature preparation step, the method is based on a simple DL network of five FCLs.

Note that for methods taking the secondary structure as input, sequence information is still represented. In RNAGCN, the nodes in the graph represent nucleotides, and edges represent pairings. In nRC and Graph+DL, the motifs that are extracted from the secondary structure are defined in relation to specific nucleotides.

We present in Fig 2 a chronological view of the methods for ncRNA classification described above. As we can see in the figure, most methods are based on CNNs, RNNs, or a combination of the two. Moreover, approaches can be separated based on the type of input data they require. A majority of DL ncRNA classifiers use the sequence as input, three use the secondary structure, and two use the sequence in combination with the structure or with epigenetic data.

**Fig 2.**
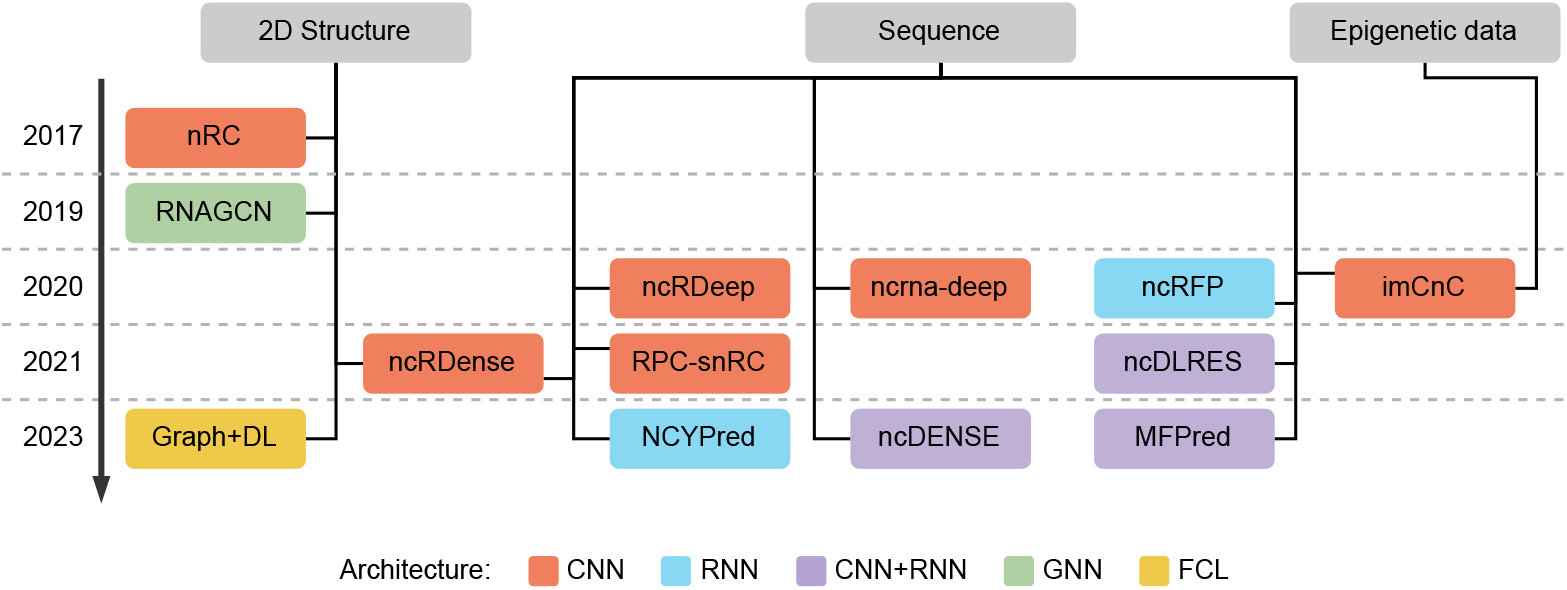
Overview of existing approaches for ncRNA classification. Methods are represented in chronological order. We specify the type of input data used (secondary structure, sequence, or epigenetic data) and the type of DL architecture (CNN, RNN, both, or other).

#### State-of-the-art datasets

Along with the tools from the previous section, multiple datasets have been proposed for evaluation of DL ncRNA classifiers in recent publications. We identify five.

The first dataset was proposed in 2017 by Fiannaca et al. [33]. Sequences were downloaded from Rfam 12 [46], and the CD-HIT tool [47] was used to remove redundant sequences. A random selection was carried out to obtain (almost-) balanced classes, with a final size of 8,920 sncRNAs spanning 13 classes. Since its proposal, this dataset has been used by all state-of-the-art methods. However, it presents a data leakage problem: 347 ncRNAs are present in both the training and the test set (representing respectively 5.5% and 13.3% of the sets).

Boukelia et al. [38] constructed a second dataset in 2020. In total, 16 sncRNA classes are represented. This dataset contains 42,000 sequences of human ncRNAs downloaded from Rfam 13 [48]. It also includes four types of epigenetic modifications (from DeepBlue [49] and HHMD [50] databases). Specifically, they obtain the positions of DNA methylation, histone methylation (20 types), histone acetylation (18 types), and binding factors and regulators related to histones. For each RNA, and for each epigenetic modification, a matrix is constructed to indicate the nearest and farthest modifications, as well as the number of modifications.

The same year, Noviello et al. [35] proposed a dataset in which ncRNAs are classified into 88 Rfam families. These families are defined as groups with RNAs that are evolutionarily conserved, have evidence for a secondary structure, and have some known function. A total of 306,016 sequences are present in the dataset.

In 2023, a dataset was proposed by Lima et al. [40]. Employing a similar protocol as Fiannaca et al. [33] but with updated data from Rfam 14.3 [51], authors were able to multiply by five the number of ncRNAs in the dataset, obtaining 45,447 ncRNAs. 13 ncRNA classes are represented. Sutanto et al. [45] also proposed a dataset in 2023. They obtained sequences from Rfam 14.1 [51] and removed those that could not be used by their structure prediction tool of choice. Then, CD-HIT [47] was used to remove redundancy in the dataset. In total, the dataset comprises 59,723 RNAs representing 12 classes.

The comparison between state-of-the-art datasets can be summed up to the origin of data, diversity of sequences, and classes represented:

- As one of the largest databases on ncRNAs, all datasets are obtained from Rfam, with recent releases containing more sequences. Boukelia et al.’s dataset [38] also integrates other databases for epigenetic data.
- In three cases ([33, 40, 45]), CD-HIT is used to remove sequences with similarity over 80%. This step restricts the size of the dataset, but only removes examples that would have been redundant while maintaining diversity. In some datasets, sequences containing degenerate nucleotides are removed ([35, 40]).
- Most datasets represent around 13 commonly-used sncRNA classes, grouping ncRNAs that share a function while not being too specific. Only Noviello et al.’s dataset [35] differs, using ncRNA families instead.

As the Rfam database is used for the construction of all datasets, it is important to note one of its limitations regarding ncRNA classes, as presented in [48]. Rfam associates classes to ncRNAs using the Infernal tool [6], which is designed to recognize sequences with high similarity. Pseudogenes, which are copies of functional genes that have mutated and become non-functional during evolution, can have high homology with the sequence they are derived from. There are no effective tools to differentiate one from the other. Therefore, some pseudogenes might be annotated as belonging to their class of origin. This is especially true in eukaryotes.

## Results

In this section, we compare the performance of different available state-of-the-art ncRNA classifiers: nRC [33], RNAGCN [44], ncrna-deep [35], MFPred [43], ncRDense [37], and NCYPred [40]. In the first subsection, their performance is measured using cross-validation and on a held-out test set on three datasets: Dataset1, Dataset1-nd and Dataset2. Dataset1 is the one by Fiannaca et al. [33], in which we fix the data leakage issue. Dataset1-nd is the same dataset in which we remove sequences containing degenerate nucleotides. Dataset2 is the one presented by Lima et al. [40] (see Supplementary Materials Table 1). In the next two subsections, where we evaluate the robustness of methods, and measure resource intensity, we focus on Dataset1-nd as it can be used by all methods. The selection of tools and datasets for this benchmark, as well as the experimental protocol, are detailed in the Methods section.

### Overall performance

In this section, performance is measured using the following multi-class metrics: Accuracy, Matthews Correlation Coefficient (MCC), F1-score, and Specificity. The formulas are detailed in the Methods section; additional scores and numerical values are presented in Supplementary Section 2.

#### 10-fold cross-validation

We performed 10-fold cross-validation with tools that could be retrained (nRC, RNAGCN, ncrna-deep, and MFPred). Results are presented in Fig 3. Scores are, in general, better on Dataset2. This can be explained by the fact that Dataset2 is much larger than Dataset1(-nd). Models are trained on a larger variety of ncRNAs, making the prediction of new examples easier. Also of note, since Dataset1-nd is very similar to Dataset1 (it is only slightly smaller due to the removal of sequences with degenerate nucleotides), the close obtained scores are expected. We notice that ncrna-deep reaches the highest scores across the different metrics and for the three datasets. MFPred and RNAGCN also obtain high scores on Dataset2, benefiting from the increased amount of data. nRC obtains lower scores across datasets, and its results vary more for the different folds compared to other tools.

**Fig 3.**
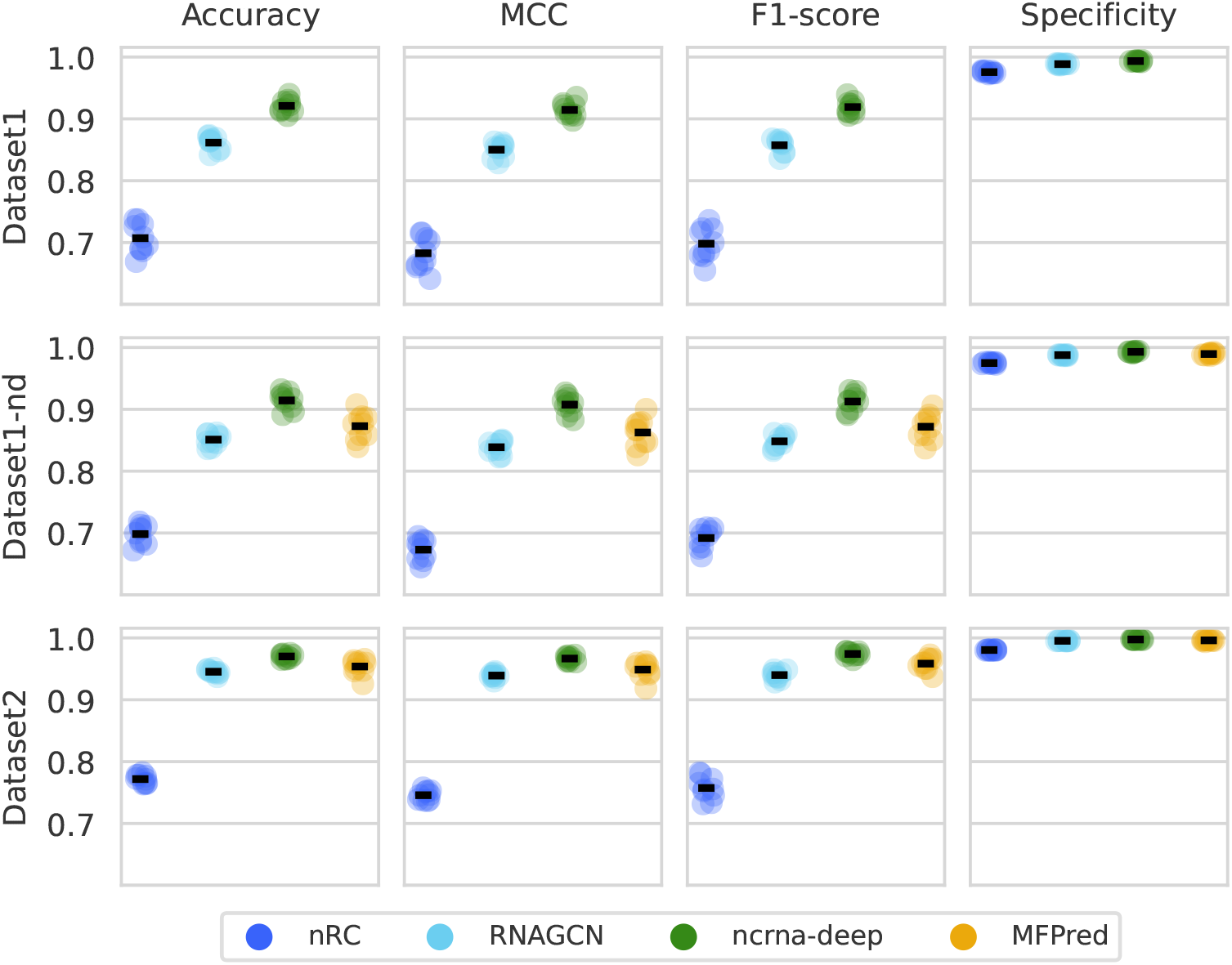
Cross-validation results. Each row corresponds to a dataset, and and each column to a metric. Each color corresponds to a method. Results on each validation set are represented by circles of the methods’ colors, on top of which a line represents the mean.

#### Performance on held-out test set

We assess in Fig 4 the prediction performance of state-of-the-art ncRNA classifiers on held-out test sets. nRC, RNAGCN, ncrna-deep and MFPred can be retrained, while ncRDense and NCYPred can only be used for prediction. This difference is visible in the results: the latter tools only perform well on the dataset they were originally trained on (Dataset1 for ncRDense and Dataset2 for NCYPred). This is particularly visible looking at the F1-score. Opposingly, tools that were retrained have consistent performances across datasets. The test scores are concordant with cross-validation results, with test scores similar to the cross-validation mean.

**Fig 4.**
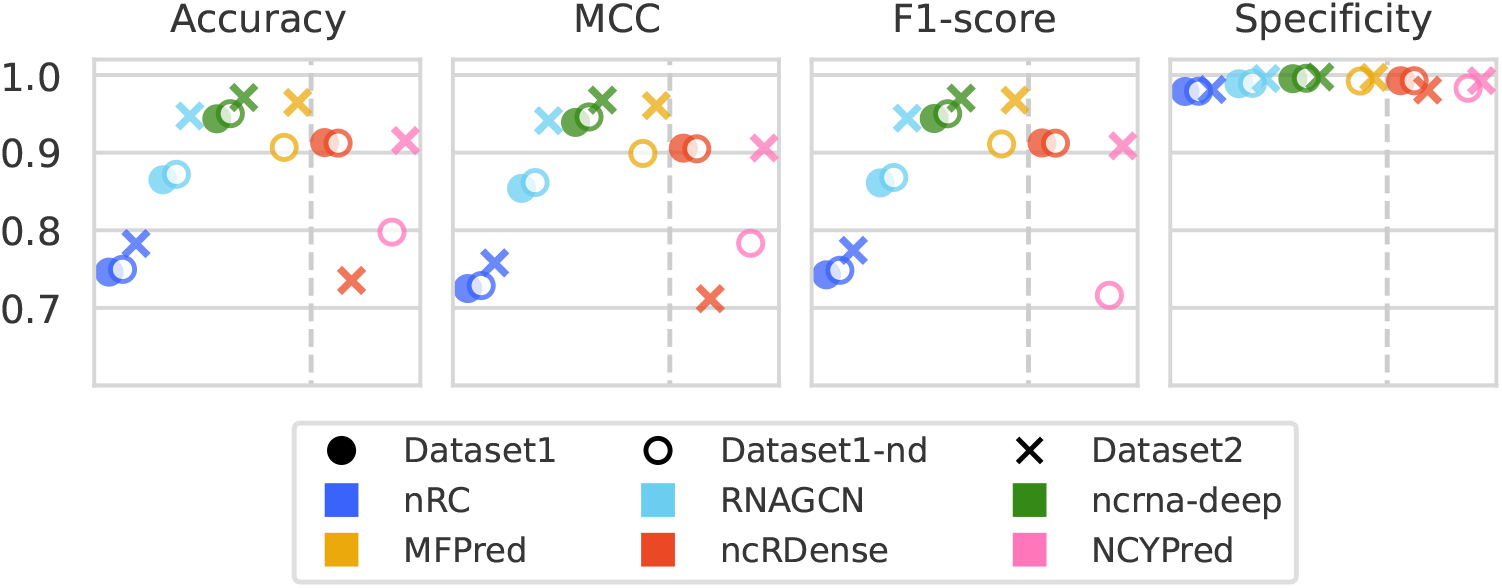
Comparison of overall performance of the state-of-the-art methods on the test sets of Dataset1, Dataset1-nd and Dataset2. The shape of the markers designates the datasets, and each color corresponds to a method. Tools on the right of the dashed line, ncRDense and NCYPred, could not be retrained before prediction. No results are shown for MFPred and NCYPred on Dataset1 since these two tools do not process degenerate nucleotides. The tools are sorted chronologically on each of the two sides separated by the dashed line.

#### Comparison with reported performance

We compare the performance we measured for each tool to the performance reported in its original publication, and illustrate the difference in Fig 5. All publications use the same formula for MCC, therefore, we base our comparison on this metric. nRC, RNAGCN, ncrna-deep, and ncRDense all use Dataset1, and MFPred and NCYPred use Dataset1-nd (albeit the biased versions with data leakage). Results in our experiments were significantly worse than what was originally reported by the various publications. These drops in performance are partially explained by the correction of the data leakage problem. For NCYPred, our lower measured performance could also be due to model training. In their evaluations, authors retrained their tool on Dataset1-nd, which we could not do, only having access to the web server version. This means that two classes from Dataset1-nd are unknown by the model and cannot be predicted. One exception is RNAGCN, for which the model we selected during hyperparameter search performs better than the one from the original publication. Only two methods, MFPred and NCYPred, reported results on Dataset2 as it is very recent. These reported results seem more reliable as we only observe negligible differences in MCC.

**Fig 5.**
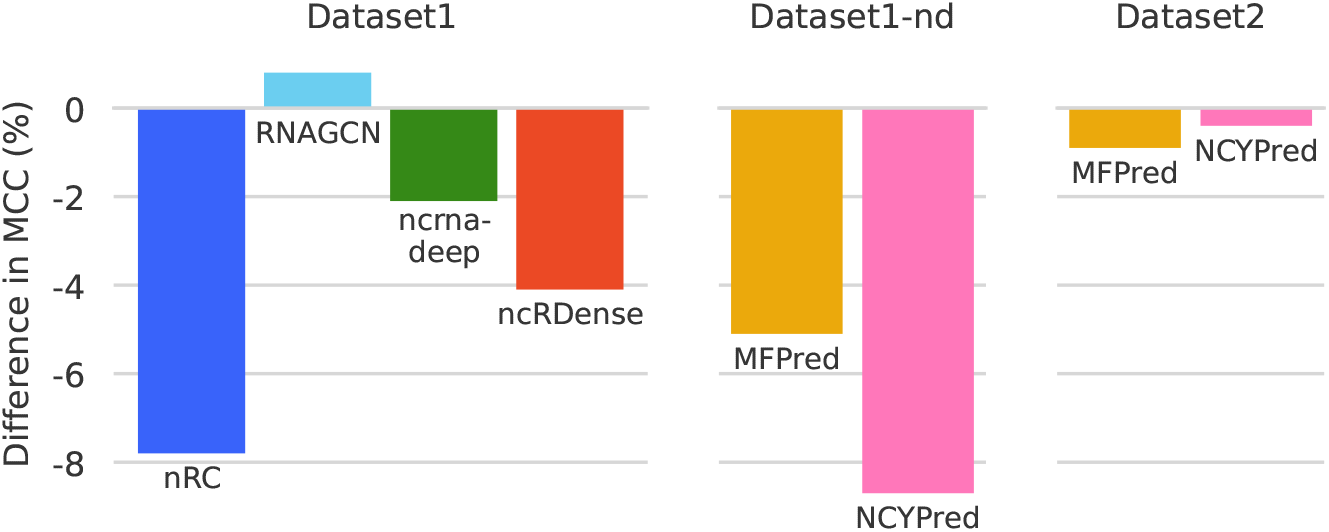
Difference between MCC values measured in our benchmark and those reported by each state-of-the-art method. A negative value of −*x* signifies that the value we measured is lower by *x* than the one originally reported.

#### Performance on individual classes

A great challenge in ncRNA classification is achieving high predictions across classes. In Fig 6, we present the accuracy of predictions for each class.

**Fig 6.**
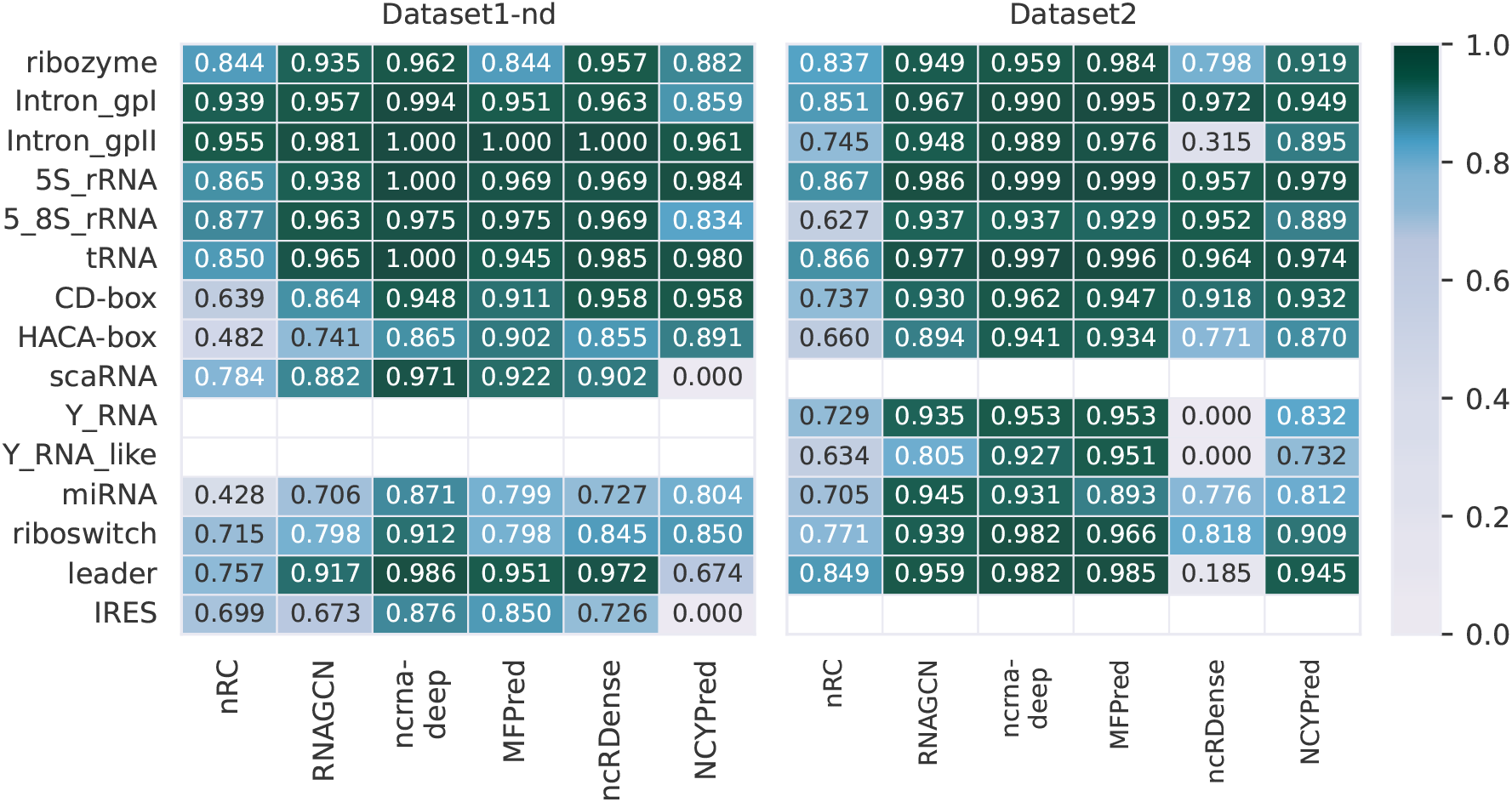
Comparison of accuracy of prediction of each ncRNA class obtained by state-of-the-art tools on Dataset1-nd and Dataset2. Light colors correspond to lower accuracies, while colors tending towards dark green represent the best results. Results for Dataset1 are comparable to those for Dataset1-nd and are presented in Supplementary Materials Fig 1.

Here, another explanation for the better results on Dataset2 than Dataset1(-nd) that were noticed in the previous section is presented: the classes included in each dataset differ, which can impact performance. The two classes included solely in Dataset1(-nd) (IRES and scaRNA) seem more difficult to predict, thus lowering overall scores. We observe once again that highest performance is obtained by ncrna-deep, seconded by MFPred. The difference between classes is striking for nRC, which is able to predict some classes quite well (Intron-gpII, 5S-rRNA), but performs poorly on other classes (miRNA, HACA-box). Aside from the classes it was not trained on (IRES and scaRNA), NCYPred obtains similar performances to other tools on Dataset1-nd. The same cannot be said for ncRDense, which does not only fail to predict the classes it was not trained on (Y RNA and Y RNA-like), but also obtains relatively poor performances for most other classes in Dataset2, underscoring the fact that not allowing to retrain a tool can be detrimental. It is interesting that, although recent tools make generally very accurate predictions, the more problematic classes are the same as those for earlier methods, such as miRNA or HACA-box. Wrongly classified miRNAs tend to be predicted as CD-box (see Supplementary Materials Figs 2-4): there is literature explaining possible similarities between the two classes [52,53]. HACA-box tends to be mispredicted as CD-box or scaRNA. These are subclasses of snoRNAs, which can explain the confusion.

### Robustness to noise

To measure the robustness of the tools to noise, we perform two experiments. In the first, we test whether models can reject non-functional sequences. In the second, we analyze the impact of sequence boundary noise.

#### Non-functional sequences

We show in Fig 7 the evolution of accuracy when adding non-functional sequences to Dataset1-nd. The ratio of non-functional to functional sequences varies from 0:1, in which case there are no non-functional sequences, to 1:1, when the size of the dataset is doubled and there are as many functional ncRNAs as non-functional. We observe that MFPred, RNAGCN and nRC’s prediction accuracy remains relatively stable as the number of non-functional sequences increases. ncrna-deep’s accuracy decreases by around 13%, but performance is still relatively satisfying. This indicates that the tools are able to recognize the characteristics of functional ncRNAs. On the other hand, ncRDense and NCYPred, which cannot be retrained, are not able to recognize non-functional sequences. Their decrease in accuracy is proportional to the number of non-functional sequences added.

**Fig 7.**
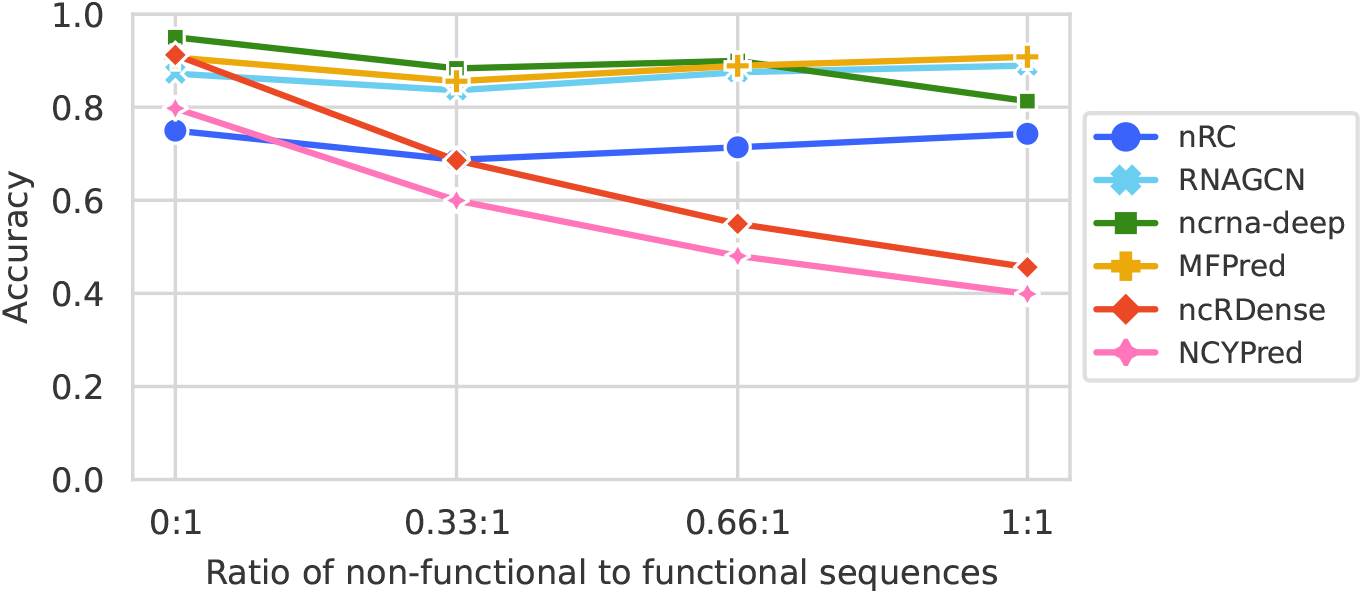
Evolution of accuracy with different numbers of non-functional sequences added to Dataset1-nd. Each line (and color) corresponds to a method. Accuracy is measured when varying the ratio of non-functional to functional sequences from 0:1 (initial dataset) to 1:1 (as many non-functional sequences as functional sequences).

#### Boundary noise

We illustrate in Fig 8 the evolution of accuracy when adding noise to Dataset1-nd sequences. We expect a drop in accuracy as the amount of noise in sequences increases and there is proportionally less information relevant to the classification task. ncrna-deep, MFPred and RNAGCN show robustness, as their accuracy only drops by around 10-15% when the noise added is equal to 200% of the sequence length. nRC is less robust, with a drop of around 25% in accuracy. ncRDense and NCYPred, which cannot be retrained, are not able to handle noisy sequences, as their accuracy drops significantly. ncRDense’s performance is particularly poor, as even with the lowest amount of noise added, its accuracy drops to around 20%.

**Fig 8.**
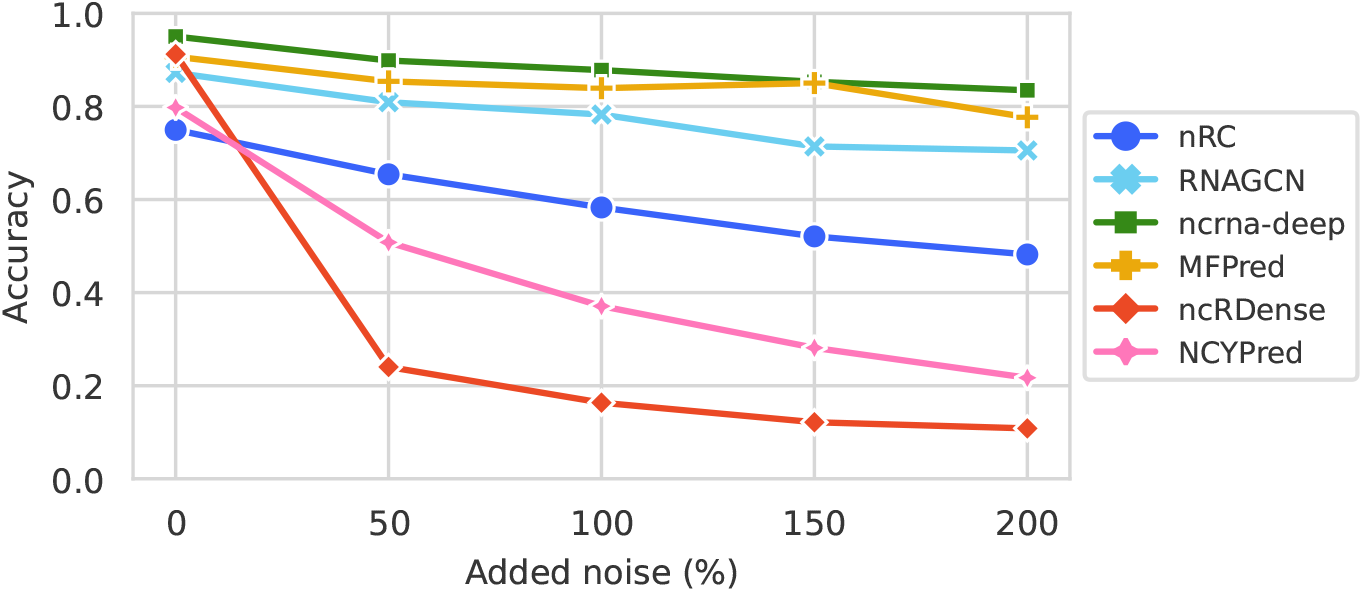
Evolution of accuracy with different percentages of added noise to Dataset1-nd sequences. Each line (and color) corresponds to a method. Accuracy is measured when the noise added to sequences is equal to 0%, 50%, 100%, 150%, or 200% of the sequence length.

### Computation time and CO_2_ emissions

We compare the computation time and CO_2_ emissions of benchmark methods, which we measured when executing the codes. RNAGCN, ncrna-deep and MFPred were executed on the same GPU machine (NVIDIA GeForce RTX 3090). nRC only supports CPU computation and was executed on Intel Xeon(R) W-1250 CPU @ 3.30GHz. CO_2_ emission estimations were conducted with CodeCarbon [54, 55]. We can use it for RNAGCN, ncrna-deep and MFPred, as source codes are all in Python, but not for nRC which is written in a mix of Java and Python.

Table 1 shows the computation time as well as the CO_2_ emissions of the state-of-art methods on Dataset1-nd. nRC and MFPred have considerably longer runtimes than RNAGCN and ncrna-deep. For nRC, this can be explained by the method’s lack of GPU support. For MFPred, the lack of speed is due to model complexity: the model is large and contains considerably more layers than the other methods. The CO_2_ emissions of these methods seem to be proportional to training time.

**Table 1.** Comparison of computation times and CO_2_ emissions on Dataset1-nd. The computation time is calculated for preprocessing, training and prediction, while the CO_2_ emission is calculated for training. Times and emissions are computed for the hyperparameter sets selected in Table 4. (See Supplementary Materials Table 4 for Dataset1 and Dataset2).

| | Computation time | | | Emissions<br>(in $gCO_2eq$) |
| --- | --- | --- | --- | --- |
|  | Preprocessing | Training | Prediction |  |
| **nRC** | 23mn 4s | 1h 57mn | 12s | - |
| **RNAGCN** | 2mn 43s | 27mn 10s | 0.4s | 9.08 |
| **ncrna-deep** | 2s | 59s | 0.6s | 0.35 |
| **MFPred** | 32s | 1h 55mn | 11s | 48.48 |

For the web servers ncRDense and NCYPred, it is difficult to measure prediction time precisely, as it depends on multiple factors: network connection, user speed, etc. Moreover, ncRDense cannot predict more than 1,000 ncRNAs at once, further complicating the measure of prediction time. As an order of magnitude, for the same 1,000 ncRNAs, we obtained predictions in around 20 seconds with NCYPred and 30 seconds with ncRDense. No preprocessing is required. CO_2_ emissions cannot be measured.

## Discussion

From the different results presented in the previous section, the current recommendation for users looking to classify ncRNAs would be to use the ncrna-deep tool, as it obtains the best results on benchmark datasets while being the fastest. MFPred, although more recent than ncrna-deep, performs slightly less well than the latter, and its execution time is longer. Moreover, it cannot be used on sequences containing degenerate nucleotides. This goes against the comparisons made in MFPred’s publication, and emphasizes the necessity of unbiased benchmarks such as the one presented here, and the importance of employing comparable experimental protocols. nRC pioneered the use of DL for ncRNA classification and was the tool of reference for years, but is now outperformed by more recent tools. RNAGCN, also obtaining lower performances, brought to the field a different point of view as its GNN-based architecture stands out from the rest of the state-of-the-art. If the goal is to classify new ncRNA datasets, we would advise against using web servers that do not provide the possibility to perform training, as it impacts performance negatively.

Several recommendations can be formulated with regard to the development of future ncRNA classifiers:

- The first choice to make in the architecture is how to represent data. nRC and RNAGCN choose to use the secondary structure. This is based on the widely accepted idea that the function of ncRNAs is linked to their structure [56], an approach adopted by many earlier ncRNA classifiers [30, 32]. It is interesting that this experimental observation does not translate particularly well to DL approaches, as methods that use the structure do not give the best results in our benchmark. This can be explained by the fact that secondary structure prediction is a difficult task, and while algorithms have improved over the years, their accuracy is still limited [57, 58]. Using newer, more accurate structure prediction algorithms could be a way to improve results in the future. Moreover, the difference in performance could also be due to the algorithms used for classification.
- Among methods that directly use the sequence, two representation approaches emerge in state-of-the-art methods: using a simple matricial representation of the sequence, or learning a meaningful representation of the sequence with RNN-based networks. These two approaches are represented in our benchmark, with ncrna-deep using a one-hot encoding of the sequence, and MFPred using four RNNs to extract different information from the sequence. Representation complexity is low for ncrna-deep, high for MFPred, but ncrna-deep performs better. The results obtained in our benchmark suggest that the simple representation should in fact be preferred, as the time gain is considerable without any effect on performance.
- While results have shown that the sequence is enough to obtain good classification results, this should not be a limit in the development of new ncRNA methods. For example, earlier ncRNA classifiers ([29, 31, 59, 60]) used RNA-seq data. The inclusion of epigenetic data by imCnC [38] also seemed promising. Indeed, while performance is an important aspect of ncRNA classification, an additional benefit in research could be to obtain a better characterization of classes – which includes describing them by patterns found at multiple levels, not just in the sequence. As stated in the introduction, the description of ncRNA classes and their functions is an ongoing research, in which computational ncRNA classifiers could be extremely useful.
- The set of ncRNA classes is bound to evolve as new ncRNA functions are discovered. On this basis, one aspect to include in future ncRNA classifiers is the ability to predict at test time classes that were unknown during training. The idea would therefore be to design a model that can identify that a sample does not belong to any of the known classes, instead of forcing classification. This could be done with zero-shot learning or abstained classification [61–63].
- Finally, there is an additional consideration regarding the availability of tools. In the description of state-of-the-art datasets, we noted an increase in dataset size and variability in the classes included. In this context, it is important to propose tools that can be applied to and evaluated on new datasets. The best practice would be to propose both a web server and to share source code. The web server would provide ease of use. By offering the possibility of retraining the model, it would not be limited to one dataset. Moreover, a well-documented source code could be used in bigger projects and with more flexibility.

## Methods

### Literature search

The scope of our study is multi-class non-coding RNA classifiers that are based on deep learning. To make sure we included all relevant literature, we performed the following search from Digital Science’s Dimensions platform (https://app.dimensions.ai): the terms *(ncRNA OR non-coding RNA) AND (deep learning) AND (classification OR classes OR families)* had to be present in the title and/or abstract. On February 14th, 2024, this search yielded 567 results. From these, a lot of results were not directly linked to our problem statement (e.g. a lot of publications were about lncRNA-disease association prediction, or cancer patient classification). We carefully checked that no relevant title had been forgotten from our study.

### Tool selection

We present in Table 2 information on the accessibility of all DL-based ncRNA classifiers. Most publications propose an online version of their tool, as a web server or by sharing source code. Web servers have the advantage of requiring no installation and being easy to use. However, they are sometimes limited: for example, here, no web server can be retrained on different datasets. If source code is available, the model can be trained on different datasets, and computational power can be increased. Nonetheless, executing the code is sometimes complicated, especially if documentation is insufficient.

**Table 2.**
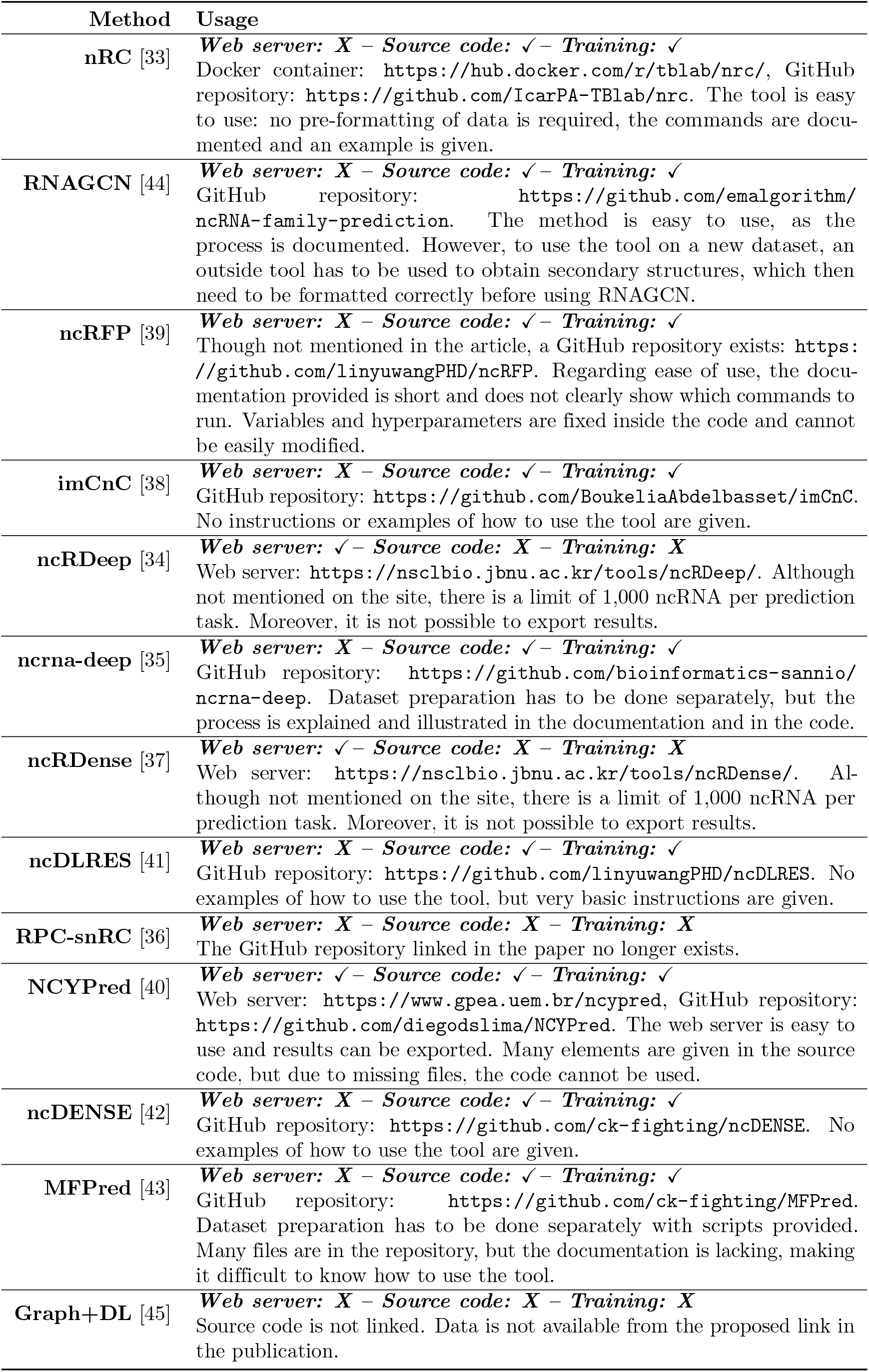
Description of the availability and ease-of-use of DL ncRNA classification tools. Tools are presented in chronological order of publication. Each cell indicates if the tool is accessible through a web server, if source code can be downloaded, and if the tool can be retrained. All links have been checked at the time of writing (February 2024).

We choose to assess the quality of prediction of six methods: nRC [33], RNAGCN [44], ncrna-deep [35], ncRDense [37], NCYPred [40] and MFPred [43]. Graph+DL [45] and RPC-snRC [36] are not available and thus cannot be tested. For a fair comparison, we need to be able to compare tools on the same datasets – this cannot be done for imCnC [38], the only tool to require epigenetic data, thus it is not included in this comparison. Finally, for tools that were developed by the same group, ample comparisons between old and new versions are already available. This concerns on one hand ncRDeep [34] and ncRDense [37], and on the other hand ncRFP [39], ncDLRES [41], ncDENSE [42] and MFPred [43]. In both cases, we use the latest version: ncRDense and MFPred.

### Dataset selection

Out of the five state-of-the-art datasets described previously, three have been made available publicly: Fiannaca et al.’s dataset [33], Noviello et al.’s dataset [35], and Lima et al.’s dataset [40]. We present in Table 3(a) information regarding the size of these datasets, as well as the number of times they were used in other publications.

**Table 3.**
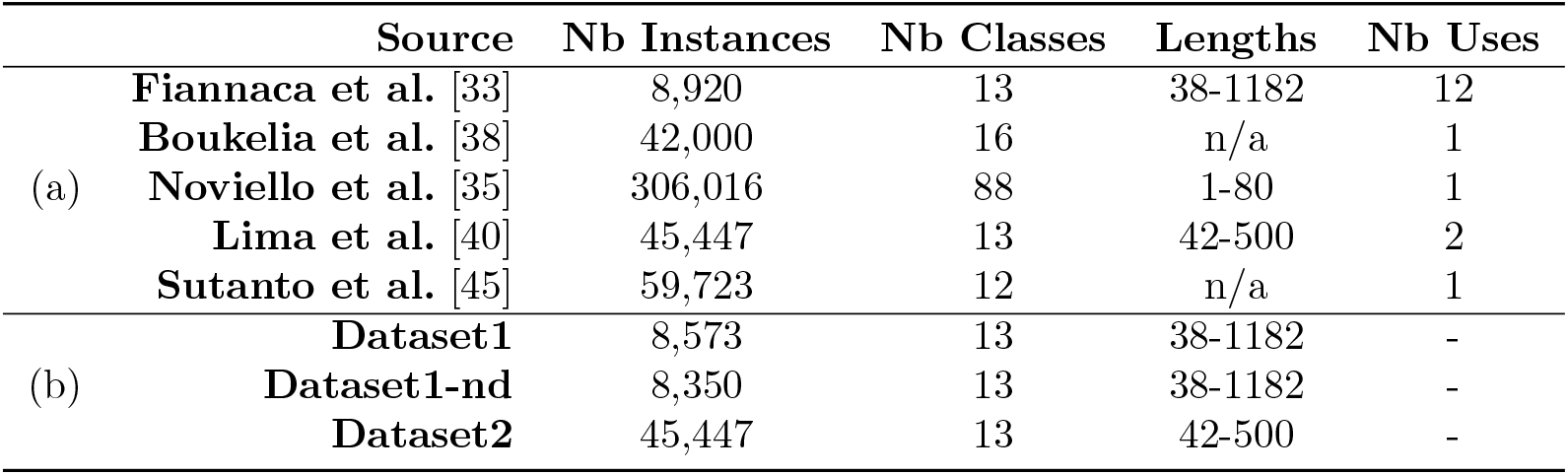
Description of datasets for ncRNA classification. **(a)** State-of-the-art datasets. **(b)** Datasets used in this study. Datasets are sorted by date. The number of instances and classes in each dataset is given. We also give the range of sequence lengths. The last column indicates how many times the dataset has been used by state-of-the-art ncRNA classifiers. For some datasets information on length range is not available (indicated with ‘n/a’) as it is not stated in the publication, and it cannot be computed as the dataset is not available.

We decide to focus on Fiannaca et al.’s dataset and Lima et al.’s dataset. Noviello et al. present their dataset with the goal of predicting ncRNA families. As such, the number of classes is high, and they can describe very precise functions. We consider these classes to be too specific for our presented problem of ncRNA classification into broader, more general functional groups (or classes). In our experiments, we refer to three datasets, also presented in Table 3(b):

- Dataset1: the dataset by Fiannaca et al. in which we fix the data leakage issue. Specifically, we remove from the test set the 347 sequences that were also present in the training set. The training and test sets contain respectively 6,320 and 2,253 ncRNAs.
- Dataset1-nd: a version of Dataset1 in which we remove sequences containing degenerate nucleotides, as these cannot be handled by some methods. The training and test sets contain respectively 6,161 and 2,189 ncRNAs.
- Dataset2: the dataset by Lima et al., in which we confirmed there was no data leakage. It does not include any sequence with degenerate nucleotides. The training and test sets contain respectively 31,801 and 13,646 ncRNAs.

### Experimental protocol

#### Evaluation metrics

Performance is measured with Accuracy, MCC (Matthews Correlation Coefficient), F1-score, and Specificity. The last two metrics are binary metrics for which we use the macro averaging generalization for multi-class problems, where scores are computed on each class separately then averaged. The four scores are defined as follows:

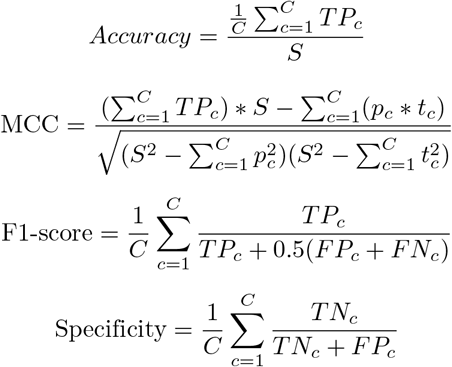

where C is the number of classes and S the number of samples. Respective of class *c, TP*_*c*_ are True Positives, *FP*_*c*_ are False Positives, *TN*_*c*_ are True Negatives and *FN*_*c*_ are False Negatives. *p*_*c*_ is the number of times class *c* was predicted, and *t*_*c*_ is the true number of members of class *c*.

#### Model training

nRC, RNAGCN, ncrna-deep and MFPred were retrained before predicting each test set. When possible, we tested different hyperparameters, which are presented in Table 4. This search could not be done for MFPred as parameter values are fixed directly in the code, and no guidelines are given on how to change them or what to change. If not stated, default hyperparameter values were used. For nRC, the model is trained for 100 epochs, with a batch size of 10. SGD optimizer is used with a learning rate of 0.05. For RNAGCN, the maximum number of epochs is set to 10,000, using the early stopping option proposed by the authors (with a patience of 100 epochs) -in general, the model stops training after 700 epochs. The batch size is 64. A learning rate of 0.0004 is used for the Adam optimizer. For ncrna-deep, 25 epochs are used, with a batch size of 32. The AMSGrad variant of the Adam optimizer is used with a learning rate of 0.0005. For MFPred, the model is trained for 100 epochs, and the batch size is set to 32. The Adam optimizer is used with a learning rate of 0.0001 and a weight decay of 0.0001. ncRDense and NCYPred were only accessible to us as web servers that could not be retrained, and were only used for prediction.

**Table 4.**
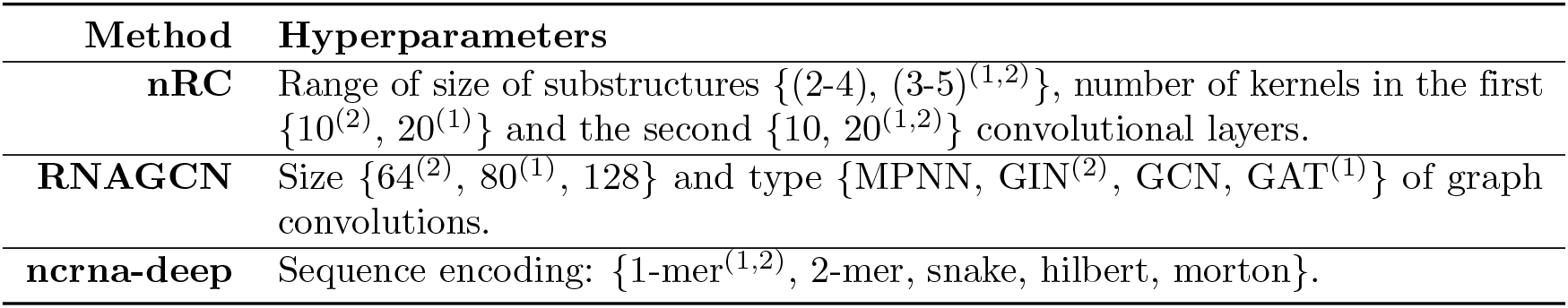
Overview of tested hyperparameters. Parameters that gave the best results on Dataset1 and Dataset2 are denoted by ^(1)^ and ^(2)^ respectively. The models chosen on Dataset1 were also used for Dataset1-nd.

#### Dataset partitionning

For 10-fold cross-validation, 10% of the training set is used as a validation set. This process is repeated 10 times, with each sample appearing exactly once in a validation set. The folds were designed in order to respect class distribution, and we ensured that there was no significant difference in terms of sequence length and presence of degenerate nucleotides. Then, models were re-trained on the full training set, and predictions were obtained on the held-out test set. MFPred and NCYPred cannot handle degenerate nucleotides, therefore their performance is not measured on Dataset1.

#### Introduction of noise

To assess whether models can reject non-functional sequences, we introduce noise by adding a new class in the dataset that contains non-functional sequences. In practice, we add to the dataset new sequences that are generated randomly, respecting di-nucleotide distribution of existing sequences. We vary the ratio of non-functional to functional sequences in *{*0:1, 0.33:1, 0.66:1, 1:1*}*. We also study the impact of sequence boundary errors by adding noise to the beginning and end of dataset sequences, while respecting mono- and di-nucleotide distribution in the sequence. The amount of noise added is proportional to sequence length: 0%, 50%, 100%, 150% and 200%. These protocols are inspired from [7, 35].

## Availability

The datasets presented in this work and the codes to reproduce this benchmark are available on the EvryRNA platform: https://evryrna.ibisc.univ-evry.fr/evryrna/ncBench.

## Footnotes

1 This method is unnamed, we refer to it using the name of its GitHub repository.

2 A k-mer is a subsequence of *k* nucleotides.

3 This method is unnamed, we refer to it by naming its two components.

## References

1. Morillon A. Non-coding RNA, Its History and Discovery Timeline; 2018. p. 1–24.

2. Lander ES. Initial impact of the sequencing of the human genome. Nature. 2011;470(7333):187–197. doi:10.1038/nature09792.

3. Kim T, Croce CM. MicroRNA: trends in clinical trials of cancer diagnosis and therapy strategies. Experimental & Molecular Medicine. 2023;55(7):1314–1321. doi:10.1038/s12276-023-01050-9.

4. Zhou Z, Wang Z, Gao J, et al. Noncoding RNA-mediated macrophage and cancer cell crosstalk in hepatocellular carcinoma. Molecular Therapy - Oncolytics. 2022;25:98–120. doi:10.1016/j.omto.2022.03.002.

5. Chen X, Luo R, Zhang Y, et al. Long noncoding RNA DIO3OS induces glycolytic-dominant metabolic reprogramming to promote aromatase inhibitor resistance in breast cancer. Nature Communications. 2022;13(1):7160. doi:10.1038/s41467-022-34702-x.

6. Nawrocki EP, Eddy SR. Infernal 1.1: 100-fold faster RNA homology searches. Bioinformatics. 2013;29(22):2933–2935. doi:10.1093/bioinformatics/btt509.

7. Navarin N, Costa F. An Efficient Graph Kernel Method for Non-Coding RNA Functional Prediction. Bioinformatics. 2017;33(17):2642–2650. doi:10.1093/bioinformatics/btx295.

8. Dupont MJ, Major F. D-ORB: A Web Server to Extract Structural Features of Related But Unaligned RNA Sequences. Journal of Molecular Biology. 2023;435(15):168181. doi:10.1016/j.jmb.2023.168181.

9. Berg MD, Brandl CJ. Transfer RNAs: Diversity in Form and Function. RNA Biology. 2021;18(3):316–339. doi:10.1080/15476286.2020.1809197.

10. Fromm B, Høye E, Domanska D, Zhong X, Aparicio-Puerta E, Ovchinnikov V, et al. MirGeneDB 2.1: Toward a Complete Sampling of All Major Animal Phyla. Nucleic Acids Research. 2021;50(D1):D204–D210. doi:10.1093/nar/gkab1101.

11. Mattick JS, Amaral PP, Carninci P, et al. Long non-coding RNAs: definitions, functions, challenges and recommendations. Nature Reviews Molecular Cell Biology. 2023; p. 1–17. doi:10.1038/s41580-022-00566-8.

12. Brayet J, Zehraoui F, Jeanson-Leh L, et al. Towards a piRNA prediction using multiple kernel fusion and support vector machine. Bioinformatics. 2014;30(17):i364–i370. doi:10.1093/bioinformatics/btu441.

13. Tran VDT, Tempel S, Zerath B, et al. miRBoost: boosting support vector machines for microRNA precursor classification. RNA. 2015;21(5):775–785. doi:10.1261/rna.043612.113.

14. Tav C, Tempel S, Poligny L, et al. miRNAFold: a web server for fast miRNA precursor prediction in genomes. Nucleic Acids Research. 2016;44(W1):W181–W184. doi:10.1093/nar/gkw459.

15. Boucheham A, Sommard V, Zehraoui F, et al. IpiRId: Integrative approach for piRNA prediction using genomic and epigenomic data. PLOS ONE. 2017;12(6):e0179787. doi:10.1371/journal.pone.0179787.

16. Baek J, Lee B, Kwon S, et al. LncRNAnet: Long non-coding RNA identification using deep learning. Bioinformatics. 2018;34(22):3889–3897. doi:10.1093/bioinformatics/bty418.

17. Liu Y, Ding Y, Li A, et al. Prediction of exosomal piRNAs based on deep learning for sequence embedding with attention mechanism. In: 2022 IEEE International Conference on Bioinformatics and Biomedicine (BIBM); 2022. p. 158–161.

18. Li M, Liang C. LncDC: a machine learning-based tool for long non-coding RNA detection from RNA-Seq data. Scientific Reports. 2022;12(1):19083. doi:10.1038/s41598-022-22082-7.

19. Raad J, Bugnon LA, Milone DH, et al. miRe2e: a full end-to-end deep model based on transformers for prediction of pre-miRNAs. Bioinformatics. 2022;38(5):1191–1197. doi:10.1093/bioinformatics/btab823.

20. Postic G, Tav C, Platon L, et al. IRSOM2: a web server for predicting bifunctional RNAs. Nucleic Acids Research. 2023;51(W1):W281–W288. doi:10.1093/nar/gkad381.

21. Rajendran V, Anandaram H, Kumar S S, et al. A Comparative Analysis of Machine Learning and Deep Learning Approaches for Circular RNA Classification. In: 2023 6th International Conference on Contemporary Computing and Informatics (IC3I). vol. 6; 2023. p. 1026–1034.

22. Krzyzanowski PM, Muro EM, Andrade-Navarro MA. Computational approaches to discovering noncoding RNA. WIREs RNA. 2012;3(4):567–579. doi:10.1002/wrna.1121.

23. Amin N, McGrath A, Chen YPP. Evaluation of deep learning in non-coding RNA classification. Nature Machine Intelligence. 2019;1(5):246–256. doi:10.1038/s42256-019-0051-2.

24. Singh D, Roy J. A large-scale benchmark study of tools for the classification of protein-coding and non-coding RNAs. Nucleic Acids Research. 2022;50(21):12094–12111. doi:10.1093/nar/gkac1092.

25. Ammunét T, Wang N, Khan S, et al. Deep learning tools are top performers in long non-coding RNA prediction. Briefings in Functional Genomics. 2022;21(3):230–241. doi:10.1093/bfgp/elab045.

26. Zhang Y, Huang H, Zhang D, et al. A Review on Recent Computational Methods for Predicting Noncoding RNAs. BioMed Research International. 2017;2017:e9139504. doi:10.1155/2017/9139504.

27. Gruber AR, Neuböck R, Hofacker IL, et al. The RNAz web server: prediction of thermodynamically stable and evolutionarily conserved RNA structures. Nucleic Acids Research. 2007;35(Web Server issue):W335–W338. doi:10.1093/nar/gkm222.

28. Lindgreen S, Gardner PP, Krogh A. MASTR: multiple alignment and structure prediction of non-coding RNAs using simulated annealing. Bioinformatics. 2007;23(24):3304–3311. doi:10.1093/bioinformatics/btm525.

29. Yuan C, Sun Y. RNA-CODE: A Noncoding RNA Classification Tool for Short Reads in NGS Data Lacking Reference Genomes. PLOS ONE. 2013;8(10):e77596. doi:10.1371/journal.pone.0077596.

30. Karklin Y, Meraz RF, Holbrook SR. Classification of non-coding RNA using graph representations of secondary structure. Pacific Symposium on Biocomputing Pacific Symposium on Biocomputing. 2005; p. 4–15.

31. Leung YY, Ryvkin P, Ungar LH, et al. CoRAL: predicting non-coding RNAs from small RNA-sequencing data. Nucleic Acids Research. 2013;41(14):e137. doi:10.1093/nar/gkt426.

32. Panwar B, Arora A, Raghava GP. Prediction and classification of ncRNAs using structural information. BMC Genomics. 2014;15(1):127. doi:10.1186/1471-2164-15-127.

33. Fiannaca A, La Rosa M, La Paglia L, et al. NRC: Non-coding RNA Classifier based on structural features. BioData Mining. 2017;10(1):27. doi:10.1186/s13040-017-0148-2.

34. Chantsalnyam T, Lim DY, Tayara H, et al. ncRDeep: Non-coding RNA classification with convolutional neural network. Computational Biology and Chemistry. 2020;88. doi:10.1016/j.compbiolchem.2020.107364.

35. Noviello TMR, Ceccarelli F, Ceccarelli M, et al. Deep learning predicts short non-coding RNA functions from only raw sequence data. PLoS Computational Biology. 2020;16(11):e1008415. doi:10.1371/journal.pcbi.1008415.

36. Asim MN, Malik MI, Zehe C, et al. A Robust and Precise ConvNet for Small Non-Coding RNA Classification (RPC-snRC). IEEE Access. 2021;9:19379–19390. doi:10.1109/ACCESS.2020.3037642.

37. Chantsalnyam T, Siraj A, Tayara H, et al. ncRDense: A novel computational approach for classification of non-coding RNA family by deep learning. Genomics. 2021;113(5):3030–3038. doi:10.1016/j.ygeno.2021.07.004.

38. Boukelia A, Boucheham A, Belguidoum M, et al. A Novel Integrative Approach for Non-coding RNA Classification Based on Deep Learning. Current Bioinformatics. 2020;15(4):338– 348. doi:10.2174/1574893614666191105160633.

39. Wang L, Zheng S, Zhang H, et al. ncRFP: A novel end-to-end method for non-coding RNAs family prediction based on Deep Learning. IEEE/ACM Transactions on Computational Biology and Bioinformatics. 2020; p. 1–1. doi:10.1109/tcbb.2020.2982873.

40. Lima DdS, Amichi LJA, Fernandez MA, et al. NCYPred: A Bidirectional LSTM Network With Attention for Y RNA and Short Non-Coding RNA Classification. IEEE/ACM Transactions on Computational Biology and Bioinformatics. 2023;20(1):557– 565. doi:10.1109/TCBB.2021.3131136.

41. Wang L, Zhong Xd, Shuo Wang, et al. ncDLRES: a novel method for non-coding RNAs family prediction based on dynamic LSTM and ResNet. BMC Bioinformatics. 2021;22(1):447. doi:10.1186/s12859-021-04365-4.

42. Chen K, Zhu X, Wang J, et al. ncDENSE: a novel computational method based on a deep learning framework for non-coding RNAs family prediction. BMC Bioinformatics. 2023;24(1):68. doi:10.1186/s12859-023-05191-6.

43. Chen K, Zhu X, Wang J, et al. MFPred: prediction of ncRNA families based on multifeature fusion. Briefings in Bioinformatics. 2023; p. bbad303. doi:10.1093/bib/bbad303.

44. Rossi E, Monti F, Bronstein M, et al. NcRNA classification with graph convolutional networks. In: Proceedings of DLG@KDD workshop. Anchorage, Alaska, US; 2019. p. 17–21.

45. Sutanto K, Turcotte M. Assessing Global-Local Secondary Structure Fingerprints to Classify RNA Sequences With Deep Learning. IEEE/ACM Transactions on Computational Biology and Bioinformatics. 2023;20(5):2736–2747. doi:10.1109/TCBB.2021.3118358.

46. Nawrocki EP, Burge SW, Bateman A, et al. Rfam 12.0: updates to the RNA families database. Nucleic Acids Research. 2015;43(Database issue):D130–D137. doi:10.1093/nar/gku1063.

47. Li W, Godzik A. Cd-hit: a fast program for clustering and comparing large sets of protein or nucleotide sequences. Bioinformatics. 2006;22(13):1658–1659. doi:10.1093/bioinformatics/btl158.

48. Kalvari I, Argasinska J, Quinones-Olvera N, et al. Rfam 13.0: shifting to a genome-centric resource for non-coding RNA families. Nucleic Acids Research. 2018;46(Database issue):D335–D342. doi:10.1093/nar/gkx1038.

49. Albrecht F, List M, Bock C, et al. DeepBlue epigenomic data server: programmatic data retrieval and analysis of epigenome region sets. Nucleic Acids Research. 2016;44(Web Server issue):W581–W586. doi:10.1093/nar/gkw211.

50. Zhang Y, Lv J, Liu H, et al. HHMD: the human histone modification database. Nucleic Acids Research. 2010;38(Database issue):D149–154. doi:10.1093/nar/gkp968.

51. Kalvari I, Nawrocki EP, Ontiveros-Palacios N, et al. Rfam 14: expanded coverage of metagenomic, viral and microRNA families. Nucleic Acids Research. 2021;49(D1):D192– D200. doi:10.1093/nar/gkaa1047.

52. Brameier M, Herwig A, Reinhardt R, et al. Human box C/D snoRNAs with miRNA like functions: expanding the range of regulatory RNAs. Nucleic Acids Research. 2011;39(2):675– 686. doi:10.1093/nar/gkq776.

53. Ono M, Scott MS, Yamada K, et al. Identification of human miRNA precursors that resemble box C/D snoRNAs. Nucleic Acids Research. 2011;39(9):3879–3891. doi:10.1093/nar/gkq1355.

54. Lacoste A, Luccioni A, Schmidt V, et al. Quantifying the carbon emissions of machine learning. Workshop on Tackling Climate Change with Machine Learning at NeurIPS 2019. 2019;.

55. Lottick K, Susai S, Friedler SA, et al. Energy Usage Reports: Environmental awareness as part of algorithmic accountability. Workshop on Tackling Climate Change with Machine Learning at NeurIPS 2019. 2019;.

56. Mortimer SA, Kidwell MA, Doudna JA. Insights into RNA structure and function from genome-wide studies. Nature Reviews Genetics. 2014;15(7):469–479. doi:10.1038/nrg3681.

57. Justyna M, Antczak M, Szachniuk M. Machine Learning for RNA 2D Structure Prediction Benchmarked on Experimental Data. Briefings in Bioinformatics. 2023;24(3):bbad153. doi:10.1093/bib/bbad153.

58. Sato K, Hamada M. Recent Trends in RNA Informatics: A Review of Machine Learning and Deep Learning for RNA Secondary Structure Prediction and RNA Drug Discovery. Briefings in Bioinformatics. 2023;24(4):bbad186. doi:10.1093/bib/bbad186.

59. Fasold M, Langenberger D, Binder H, et al. DARIO: a ncRNA detection and analysis tool for next-generation sequencing experiments. Nucleic Acids Research. 2011;39(Web Server issue):W112–117. doi:10.1093/nar/gkr357.

60. Videm P, Rose D, Costa F, et al. BlockClust: efficient clustering and classification of non-coding RNAs from short read RNA-seq profiles. Bioinformatics. 2014;30(12):i274–i282. doi:10.1093/bioinformatics/btu270.

61. Pourpanah F, Abdar M, Luo Y, Zhou X, Wang R, Lim CP, et al. A Review of Generalized Zero-Shot Learning Methods. IEEE Transactions on Pattern Analysis and Machine Intelligence. 2022; p. p1–20. doi:10.1109/TPAMI.2022.3191696.

62. Creux C, Zehraoui F, Hanczar B, et al. A3SOM, abstained explainable semi-supervised neural network based on self-organizing map. PLOS ONE. 2023;18(5):e0286137. doi:10.1371/journal.pone.0286137.

63. Hendrickx K, Perini L, Van Der Plas D, Meert W, Davis J. Machine Learning with a Reject Option: A Survey. Machine Learning. 2024;113(5):3073–3110. doi:10.1007/s10994-024-06534-x.

