## Supplementary Materials for "Comparison and benchmark of deep learning methods for non-coding RNA classification"

#### 1 Description of datasets

We present in Table 1 the class composition and average sequence length for Dataset1 and Dataset2.

| | **Dataset1** | | | **Dataset2** | | |
|---|---|---|---|---|---|---|
| **Label** | **Avg. length** | **Train size** | **Test size** | **Avg. length** | **Train size** | **Test size** |
| **5S rRNA** | 119.1 | 500 (11) | 197 (5) | 119.9 | 3,496 (0) | 1,500 (0) |
| **5.8S rRNA** | 153.4 | 500 (58) | 184 (21) | 150.7 | 322 (0) | 126 (0) |
| **CD-box** | 106.0 | 500 (4) | 195 (4) | 108.3 | 3,492 (0) | 1,504 (0) |
| **HACA-box** | 139.8 | 500 (2) | 194 (1) | 144.7 | 3,514 (0) | 1,485 (0) |
| **Intron gpI** | 342.5 | 500 (46) | 174 (11) | 284.5 | 903 (0) | 390 (0) |
| **Intron gpII** | 95.6 | 500 (12) | 160 (5) | 139.8 | 2,475 (0) | 1,089 (0) |
| **IRES** | 233.5 | 320 (8) | 124 (11) | - | - | - |
| **leader** | 124.7 | 500 (3) | 145 (1) | 212.9 | 3,481 (0) | 1,514 (0) |
| **miRNA** | 108.5 | 500 (4) | 196 (2) | 116.4 | 3,529 (0) | 1,466 (0) |
| **riboswitch** | 142.1 | 500 (2) | 193 (0) | 142.2 | 3,512 (0) | 1,483 (0) |
| **ribozyme** | 259.8 | 500 (4) | 188 (2) | 303.1 | 3,218 (0) | 1,408 (0) |
| **scaRNA** | 174.2 | 500 (2) | 103 (1) | - | - | - |
| **tRNA** | 77.7 | 500 (3) | 200 (0) | 81.4 | 3,463 (0) | 1,533 (0) |
| **Y RNA** | - | - | - | 104.7 | 320 (0) | 107 (0) |
| **Y RNA-like** | - | - | - | 129.3 | 76 (0) | 41 (0) |
| **ALL** | 157.3 | 6,320 (159) | 2,253 (64) | 154.5 | 31,801 (0) | 13,646 (0) |

**Table 1. Description of ncRNA classification datasets. Dataset1**: a version of the dataset in Fiannaca et al. in which we removed the data leakage bias. **Dataset2**: the dataset by Lima et al. We present the average length of sequences from each class, as well as the number of ncRNAs in the training and test sets. In parenthesis is indicated the number of sequences containing degenerate nucleotides (i.e., other than A,C,G,T/U). These are removed from Dataset1 to form Dataset1-nd.

#### 2 Additional results

We present in Table 2 the numerical results of cross-validation. These scores correspond to Fig 3 in the main manuscript.

| | Method | Accuracy | MCC | F1-score |
|---|---|---|---|---|
| **Dataset1** | **nRC** | 0.707 ± 0.024 | 0.683 ± 0.026 | 0.698 ± 0.026 |
| | **RNAGCN** | 0.862 ± 0.011 | 0.85 ± 0.012 | 0.857 ± 0.011 |
| | **ncrna-deep** | 0.921 ± 0.01 | 0.914 ± 0.011 | 0.919 ± 0.01 |
| **Dataset1-nd** | **nRC** | 0.698 ± 0.015 | 0.673 ± 0.017 | 0.692 ± 0.016 |
| | **RNAGCN** | 0.851 ± 0.01 | 0.839 ± 0.01 | 0.848 ± 0.01 |
| | **ncrna-deep** | 0.914 ± 0.013 | 0.908 ± 0.014 | 0.913 ± 0.014 |
| | **MFPred** | 0.873 ± 0.021 | 0.863 ± 0.022 | 0.872 ± 0.021 |
| **Dataset2** | **nRC** | 0.772 ± 0.007 | 0.746 ± 0.008 | 0.757 ± 0.018 |
| | **RNAGCN** | 0.945 ± 0.004 | 0.939 ± 0.004 | 0.94 ± 0.007 |
| | **ncrna-deep** | 0.97 ± 0.004 | 0.967 ± 0.004 | 0.974 ± 0.004 |
| | **MFPred** | 0.954 ± 0.012 | 0.949 ± 0.013 | 0.958 ± 0.01 |

| | Method | Recall | Precision | Specificity |
|---|---|---|---|---|
| **Dataset1** | **nRC** | 0.7 ± 0.025 | 0.7 ± 0.026 | 0.976 ± 0.002 |
| | **RNAGCN** | 0.857 ± 0.011 | 0.861 ± 0.012 | 0.988 ± 0.001 |
| | **ncrna-deep** | 0.919 ± 0.011 | 0.922 ± 0.009 | 0.993 ± 0.001 |
| **Dataset1-nd** | **nRC** | 0.694 ± 0.016 | 0.694 ± 0.015 | 0.975 ± 0.001 |
| | **RNAGCN** | 0.847 ± 0.01 | 0.854 ± 0.011 | 0.988 ± 0.001 |
| | **ncrna-deep** | 0.912 ± 0.014 | 0.918 ± 0.012 | 0.993 ± 0.001 |
| | **MFPred** | 0.871 ± 0.021 | 0.879 ± 0.02 | 0.989 ± 0.002 |
| **Dataset2** | **nRC** | 0.742 ± 0.024 | 0.785 ± 0.012 | 0.98 ± 0.001 |
| | **RNAGCN** | 0.94 ± 0.012 | 0.942 ± 0.009 | 0.995 ± 0.0 |
| | **ncrna-deep** | 0.971 ± 0.007 | 0.978 ± 0.002 | 0.997 ± 0.0 |
| | **MFPred** | 0.955 ± 0.009 | 0.964 ± 0.011 | 0.996 ± 0.001 |

**Table 2. 10-fold cross-validation mean and standard deviation of different metrics.**

We present in Table 3 the numerical results of evaluation on test sets. These scores correspond to Fig 4 in the main manuscript.

| | Method | Accuracy | MCC | F1-score | Recall | Precision | Specificity |
|---|---|---|---|---|---|---|---|
| **Dataset1** | **nRC** | 0.746 | 0.725 | 0.743 | 0.753 | 0.738 | 0.979 |
| | **RNAGCN** | 0.865 | 0.854 | 0.861 | 0.864 | 0.863 | 0.989 |
| | **ncrna-deep** | 0.944 | 0.939 | 0.944 | 0.943 | 0.946 | 0.995 |
| | **ncRDense** | 0.913 | 0.906 | 0.912 | 0.91 | 0.92 | 0.993 |
| **Dataset1-nd** | **nRC** | 0.75 | 0.729 | 0.748 | 0.756 | 0.747 | 0.979 |
| | **RNAGCN** | 0.872 | 0.861 | 0.868 | 0.871 | 0.871 | 0.989 |
| | **ncrna-deep** | 0.95 | 0.946 | 0.95 | 0.951 | 0.951 | 0.996 |
| | **MFPred** | 0.907 | 0.899 | 0.911 | 0.909 | 0.916 | 0.992 |
| | **ncRDense** | 0.912 | 0.905 | 0.912 | 0.91 | 0.92 | 0.993 |
| | **NCYPred** | 0.798 | 0.783 | 0.716 | 0.744 | 0.705 | 0.983 |
| **Dataset2** | **nRC** | 0.783 | 0.758 | 0.774 | 0.76 | 0.793 | 0.981 |
| | **RNAGCN** | 0.947 | 0.941 | 0.944 | 0.936 | 0.954 | 0.995 |
| | **ncrna-deep** | 0.971 | 0.968 | 0.97 | 0.965 | 0.976 | 0.998 |
| | **MFPred** | 0.965 | 0.961 | 0.968 | 0.962 | 0.974 | 0.997 |
| | **ncRDense** | 0.735 | 0.712 | 0.52 | 0.562 | 0.549 | 0.981 |
| | **NCYPred** | 0.916 | 0.906 | 0.909 | 0.895 | 0.926 | 0.993 |

**Table 3. Evaluation of different metrics on held-out test sets.**

In the main manuscript, per-class accuracy is presented on Dataset1-nd and Dataset2. Dataset1 and Dataset1-nd are very similar, therefore, results do not change much. This is proven by Figure 1, where we present per-class accuracy on Dataset1, and once again on Dataset1-nd for easier comparison.

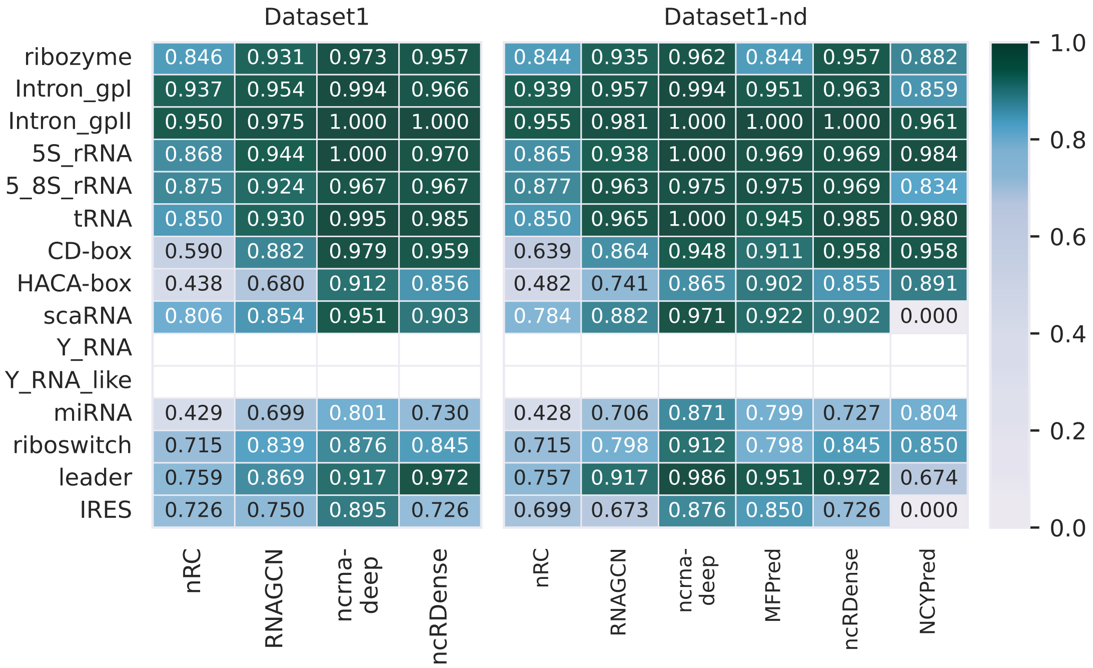


**Fig 1. Comparison of accuracy of prediction of each ncRNA class obtained by state-of-the-art tools on Dataset1 and Dataset1-nd.** Light colors correspond to lower accuracies, while colors tending towards dark green represent the best results. Note that results cannot be obtained on Dataset1 for MFPred and NCYPred as these methods cannot predict sequences containing degenerate nucleotides.

#### 3 Computation time and $CO_2$ emissions

In Table 4, computation times and $CO_2$ emissions are presented for Dataset1 and Dataset2.

| | | Computation time | | | Emissions |
|---|---|---|---|---|---|
| | **Preprocessing** | **Training** | **Prediction** | (in $gCO_2eq$) | |
| **Dataset1** | **nRC** | 23mn 54s | 2h | 12s | - |
| | **RNAGCN** | 2mn52 | 17mn 17s | $< 1s$ | 5.9 |
| | **ncrna-deep** | 2s | 59s | $< 1s$ | 0.3 |
| **Dataset2** | **nRC** | 3h 7mn | 9h 4mn | 57s | - |
| | **RNAGCN** | 13mn 22s | 1h 5mn | 2s | 17.8 |
| | **ncrna-deep** | 11s | 4mn 11s | 2s | 1.7 |
| | **MFPred** | 2mn 53s | 9h 19mn | 1mn 3s | 233.8 |

**Table 4. Comparison of computation times and $CO_2$ emissions on Dataset1 and Dataset2.** The computation time is calculated for preprocessing, training and prediction, while the $CO_2$ emission is calculated for training.

### 4 Confusion matrices

In this section, we present the normalized confusion matrices for all benchmark methods. This is an extension of the results presented in Figure 5 in the main manuscript and in Figure 1.

Normalized confusion matrices for the tests sets of Dataset1, Dataset1-nd and Dataset2 are presented, respectively, in Figure 2, Figure 3, and Figure 4.

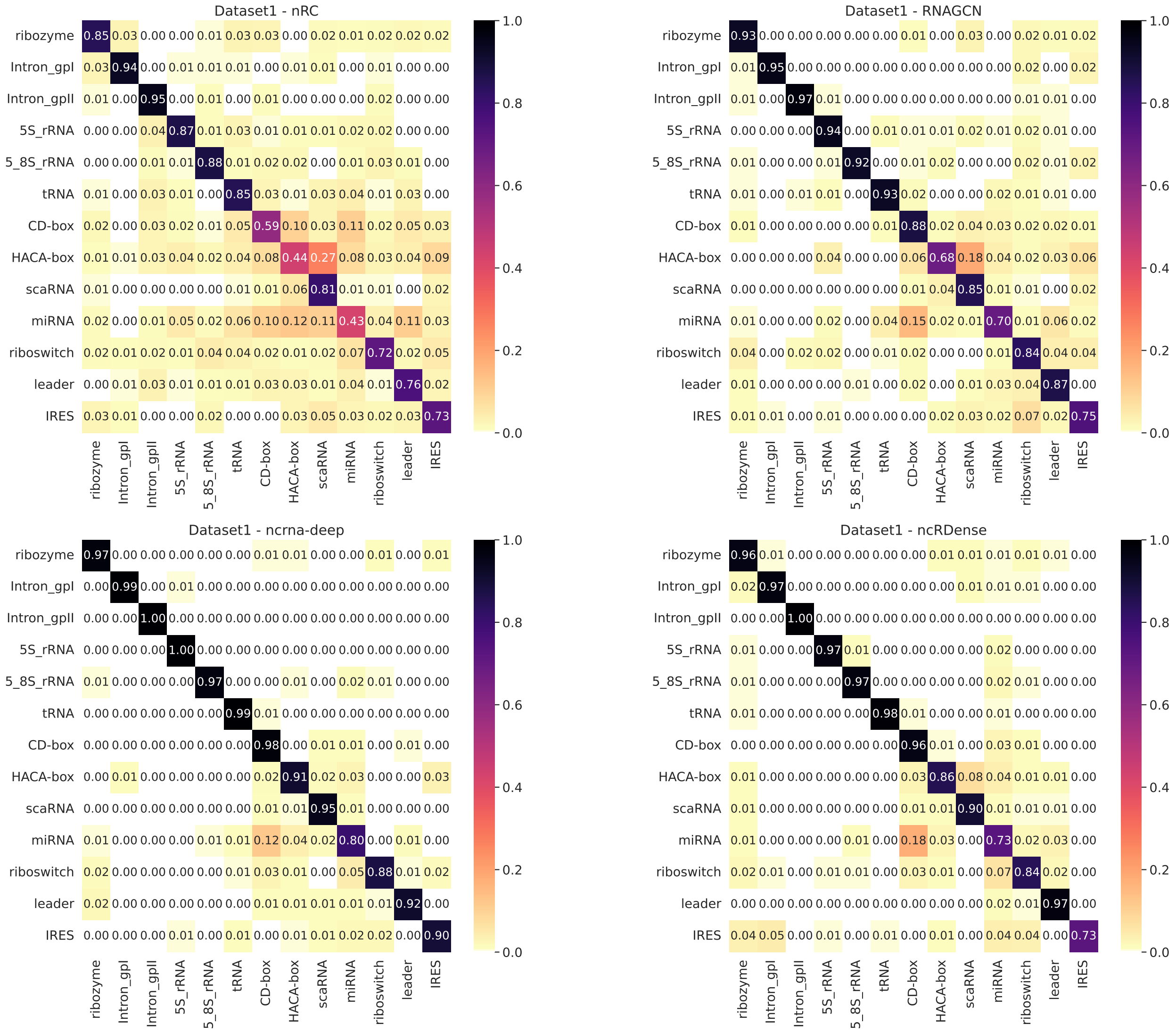


**Fig 2. Confusion matrices on Dataset1.**

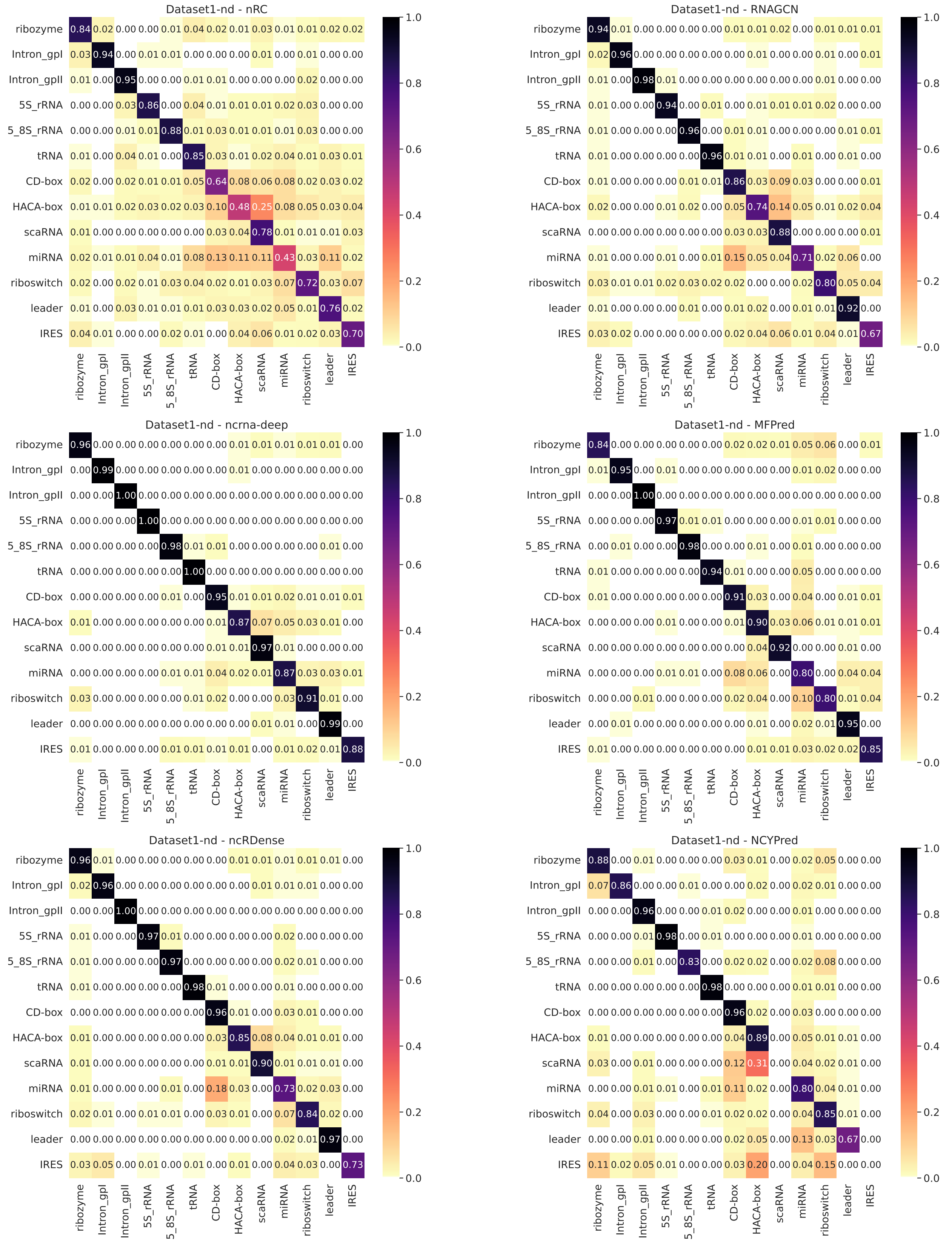


**Fig 3. Confusion matrices on Dataset1-nd.**

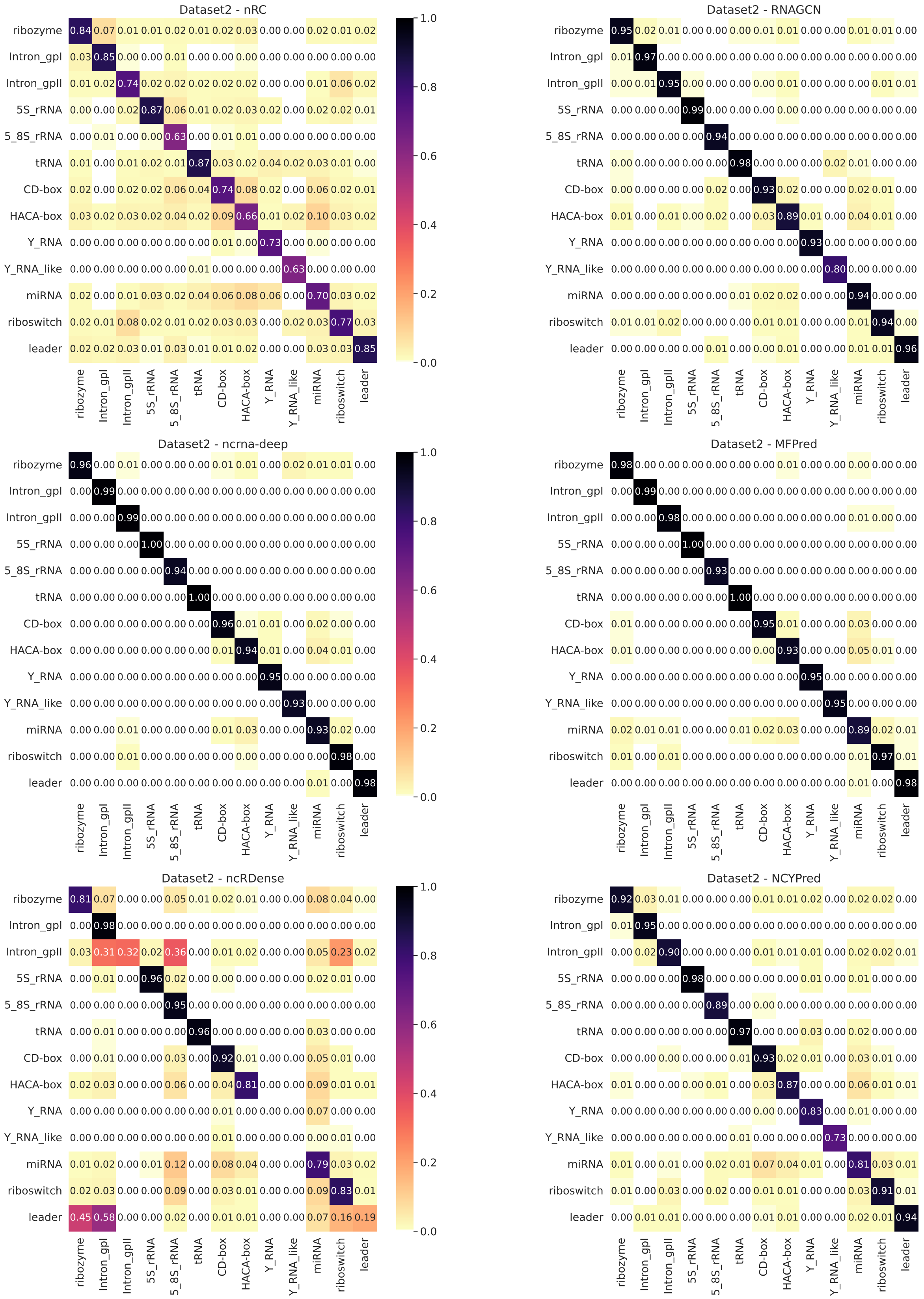


**Fig 4. Confusion matrices on Dataset2.**
